# Pathology-targeted EP4 agonism resolves fibrosis and restores regeneration in a rat model of adolescent Duchenne muscular dystrophy

**DOI:** 10.64898/2026.05.29.728674

**Authors:** Nasim Kajabadi, Ashok Narasimhan, Gang Chen, Lin Yi, Cristina Rodríguez-Rodríguez, Ian F. Coccimiglio, Tiffany Huang, Mariana Rendeiro, Keitaro Yamanouchi, Urs O. Häfeli, Paul Kostenuik, Robert N. Young, Fabio M.V. Rossi

**Affiliations:** University of British Columbia; UBC; SFU; University of Tokyo; University of Michigan School of Dentistry

## Abstract

Duchenne muscular dystrophy (DMD) presents a critical therapeutic gap in adolescent patients, where extensive fibro-fatty muscle replacement and depletion of the regenerative niche render existing interventions insufficient. Prostaglandin E2 signaling through the EP4 receptor stimulates bone and muscle regeneration and repair, but systemic off-target effects have limited the clinical translation of EP4 agonism in diseases such as DMD. We therefore evaluated irodanoprost (IROD), a bone-targeted prodrug of an EP4-selective agonist, in a DMD rat model, comparing early- and late-intervention cohorts. In adolescent rats, 8 weeks of treatment reduced body weight deficit by 41.4% and restored hindlimb muscle mass and maximum tetanic force to wild-type levels. IROD dose-dependently inhibited fibro-adipogenic progenitor differentiation into α-SMA⁺ myofibroblasts, facilitating active resolution of established fibrosis below pre-treatment baseline. This was accompanied by re-activation of a synchronized regenerative program marked by clustered eMHC⁺ fibers, restoring the total myofiber pool to wild-type levels. A strong linear correlation between intramuscular fat reduction and fibrosis resolution suggests that MRI-based fat imaging may serve as a non-invasive surrogate for monitoring anti-fibrotic efficacy. The efficacy of IROD is likely aided by the fact that while it selectively distributes to bone in healthy animals, we observed markedly enhanced accumulation of the drug in dystrophic muscle. These findings establish IROD as a pathology-targeted approach capable of resolving the fibro-fatty niche and restoring regenerative capacity in advanced DMD.

**Summary:** Pathology-targeted EP4 agonism resolves established fibrosis and restores myofiber regeneration in a rat model of adolescent Duchenne dystrophy.

**Graphic abstract:** 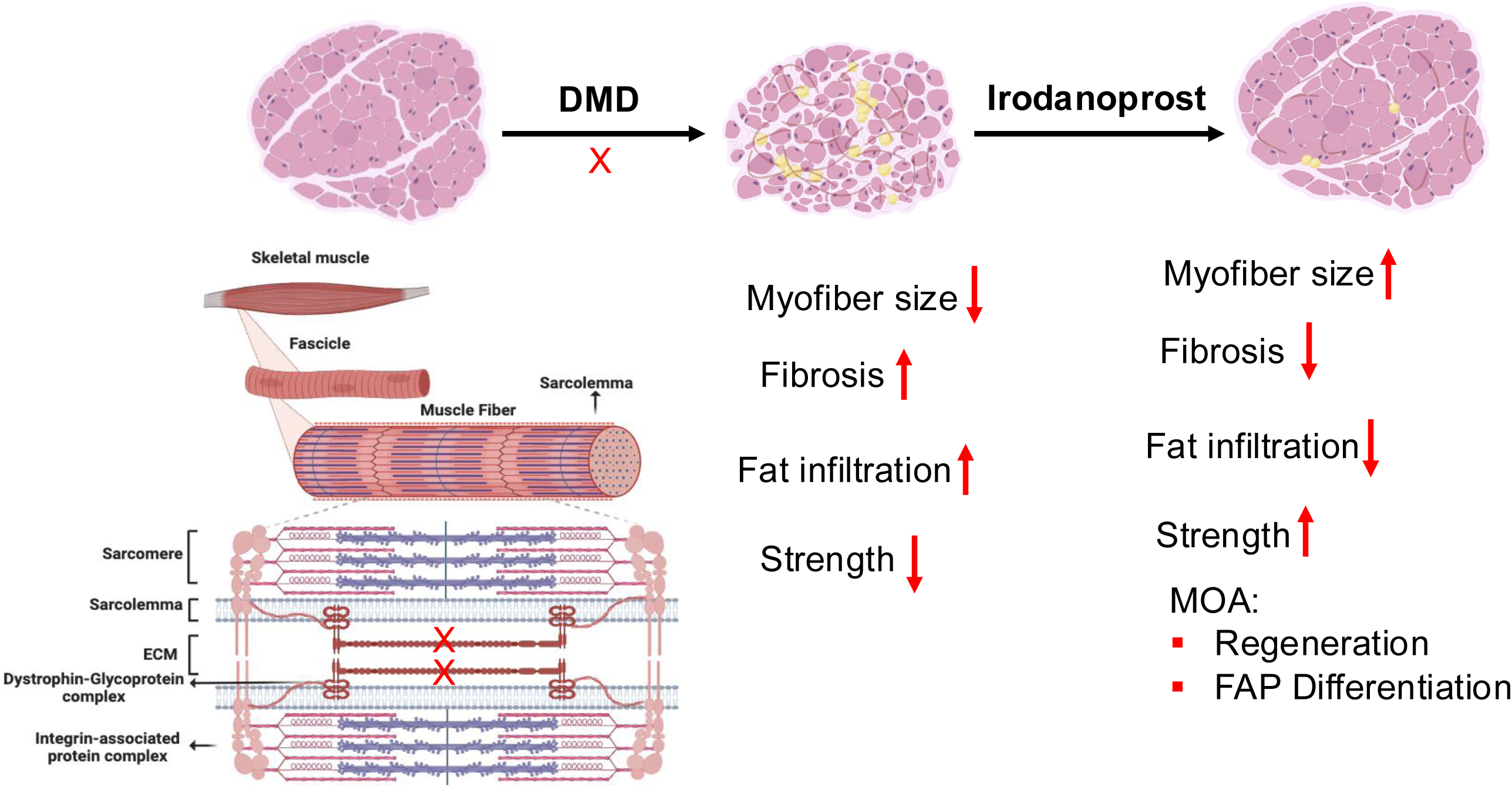

## Introduction

Duchenne muscular dystrophy (DMD) is among the most severe inherited disorders of childhood. Caused by loss-of-function mutations in the *DMD* gene, the complete absence of the cytoskeletal protein dystrophin renders myofibers mechanically fragile, initiating relentless cycles of degeneration that progressively consume the musculature, abolish ambulation, and ultimately cause premature death from cardiorespiratory failure(*1, 2*). Despite decades of research, a fundamental gap persists between the promise of emerging therapies and their reach in clinical practice. Exon-skipping oligonucleotides, micro-dystrophin gene replacement, and cell-based approaches have advanced the field considerably, yet nearly all are designed and trialed exclusively in young children aged 4–7 years, when sufficient residual muscle remains to benefit from genetic correction(*1, 3, 4*). Adolescent and adult patients, who constitute a large and growing fraction of the DMD population, are largely excluded from these interventions. By the time they reach their teenage years, extensive fibro-fatty replacement, chronic inflammation, and progressive depletion of the muscle satellite cell (MuSC) pool have fundamentally altered the biological landscape, rendering it hostile to repair and refractory to gene-based stabilization strategies(*1, 5*).

The central biological barrier to treating advanced DMD is the collapse of the regenerative niche. In early disease, MuSCs mount a vigorous but ultimately unsustainable compensatory response to myofiber loss. Repeated cycles of degeneration gradually exhaust this pool and promote the pathological expansion of fibro-adipogenic progenitors (FAPs)—a stromal PDGFRα+ population that, under TGF-β–driven fibrogenic cues, differentiates into myofibroblasts and deposits collagen throughout the interstitium(*6, 7*). This fibro-fatty matrix simultaneously replaces functional contractile tissue and creates an inhibitory niche that suppresses residual regenerative activity (*8*). Current standard-of-care corticosteroids delay disease progression but do not reverse fibrosis, restore myofiber number, or reactivate a stalled myogenic program, and are associated with significant adverse effects including mood alterations, weight gain, osteoporosis and fragility fractures with long-term use(*1, 9*). No approved therapy is capable of resolving established pathology and re-engaging productive muscle regeneration in adolescent DMD. Prostaglandin E2 (PGE2) signaling through the EP4 receptor has emerged as a critical endogenous regulator of bone anabolism (*10*) as well asMuSC expansion and muscle regeneration following injury. EP4 activation elevates intracellular cAMP, promotes satellite-cell entry into the cell cycle via a cAMP/phospho-CREB/Nurr1 transcriptional axis, and accelerates muscle repair in young and aged preclinical models.(*11, 12*). However, systemic EP4 agonism is constrained by cardiovascular and gastrointestinal off-target effects—including hypotension and colonic hyperemia—that arise from the ubiquitous expression of EP4 across vascular, cardiac, and mucosal tissues, limiting clinical translation(*13, 14*). Tissue-targeted delivery of EP4 agonism selectively to the dystrophic niche has therefore remained an unmet therapeutic challenge.

To address this, we evaluated Irodanoprost (IROD), a bone-targeted prodrug comprising a bis-phosphonate-based bone-binding moiety, a hydrolyzable ester linker, and a potent EP4-selective PGE2 analog(*15*). Although originally designed to primarily target the severe osteopenia characteristic of DMD, we hypothesized that IROD could nonetheless release its highly potent and tissue-permeable EP4 agonist moiety from the bone depot and thereby affect adjacent muscles. We further considered that the increased permeability of damaged muscle to small molecules might enhance IROD exposure at sites of active pathologyWe tested this hypotheses in the DMD rat, a CRISPR/Cas9-engineered model carrying a frameshift Dmd mutation that more accurately recapitulates the severe human adolescent phenotype than the standard mdx mouse, including progressive weight loss, limb-muscle wasting, and multi-organ fibrosis(*16*). By comparing early-intervention (2-month-old) and late-intervention (7-month-old) cohorts, we asked whether targeted EP4 activation could overcome regenerative exhaustion and reverse established structural pathology at a stage currently beyond the reach of any approved or investigational therapy. We show that IROD restores myofiber number to wild-type levels, actively resolves established fibrosis and intramuscular fat, and reactivates a synchronized regenerative program that is otherwise extinguished by chronic disease—establishing a pathology-targeted framework in which disease severity itself enhances drug delivery and efficacy.

## Results

### IROD treatment produces dose-dependent rescue of muscle mass and contractile function in adolescent DMD rats

To evaluate the effects of IROD in the context of DMD pathology, we administered the drug to DMD rats, a CRISPR/Cas9-engineered model carrying an out-of-frame *Dmd* frameshift mutation that leads to the complete absence of Dystrophin and more accurately recapitulates severe human phenotypes than the *mdx* mouse(*16*). In accordance with its bone-targeting design, IROD-treated rats showed reversal of trabecular bone loss compared with untreated DMD controls, which will be described in detail elsewhere. Additionally, IROD treatment produced attenuation of progressive body weight loss, with treated DMD cohorts maintaining growth curves significantly above those of untreated DMD controls throughout the study period (Supp. Fig. 1A). To evaluate the effects of IROD at increasing doses, we treated 7-month-old DMD rats, an age developmentally equivalent to human adolescence (*17*), with weekly subcutaneous injections of vehicle (0 mg/kg), 1 mg/kg, or 3 mg/kg IROD for 8 weeks (Fig. 1A-B). Vehicle-treated DMD controls exhibited progressive body weight loss, reaching a final body weight of 412.9 ± 45.1 g compared with 594.3 ± 43.6 g in WT controls—a 30.5% deficit. IROD treatment dose-dependently attenuated this deficit: the 1 mg/kg group reached 455.3 ± 14.5 g (23.4% smaller deficit vs. WT than vehicle-treated DMD controls) and the 3 mg/kg group reached 488.0 ± 27.5 g (41.4% smaller deficit), with the 3 mg/kg effect reaching statistical significance (Fig. 1C).

**Fig. 1.**
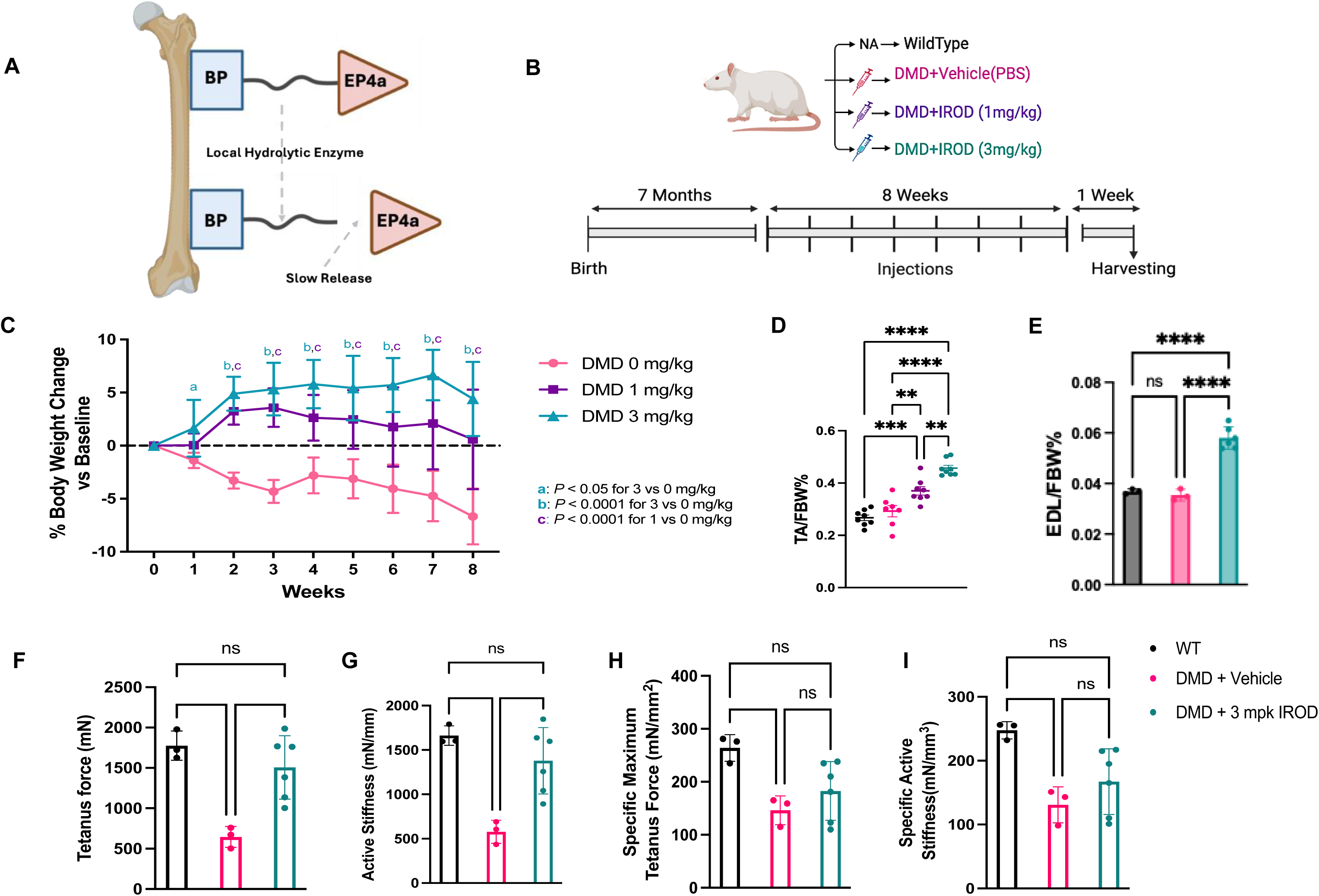
Dose-dependent systemic rescue and restoration of hindlimb muscle mass and contractile function in IROD-treated DMD rats. (A) Schematic of the IROD prodrug architecture, comprising a phosphonate-based mineral-targeting moiety, a hydrolyzable ester linker, and the EP4-selective prostaglandin E2 analog MES1002 (B) Schematic of the dose-escalation study design. Seven-month-old DMD rats received weekly subcutaneous injections of vehicle (0 mg/kg), 1 mg/kg, or 3 mg/kg IROD for 8 weeks alongside age-matched WT controls. (C) Percentage change in body weight from baseline. n = 7–8 per group. (D) Tibialis anterior (TA) muscle weight normalized to final body weight (% TA/FBW). n = 7–8 per group. (E) EDL muscle weight normalized to final body weight (% EDL/FBW). n = 3–6 per group. (F) Absolute maximum tetanic force of isolated EDL muscles. (G) Absolute active stiffness of isolated EDL muscles. (H) Specific tetanic force (absolute force normalized to muscle cross-sectional area, mCSA). No significant difference was detected between IROD-treated and untreated DMD groups after normalization. (I) Normalized active stiffness (absolute stiffness normalized to mCSA). No significant difference was detected between IROD-treated and untreated DMD groups after normalization, consistent with functional rescue driven by increased muscle mass rather than qualitative improvement of individual myofibers. For panels F–I, data are mean ± SEM; n = 7–8 per group. All statistical comparisons by two-way ANOVA; *P < 0.05, **P < 0.01, ***P < 0.001.

This systemic recovery was underpinned by significant increases in individual muscle mass across the hindlimb. Untreated DMD rats exhibited significant wasting of the tibialis anterior (TA), whereas both IROD doses significantly increased TA weight relative to vehicle-treated controls. The 3 mg/kg group reached TA weights significantly exceeding those of WT controls, an effect that remained significant after normalization to final body weight (Fig. 1D). A parallel increase in extensor digitorum longus (EDL) mass was observed at 3 mg/kg (Fig. 1E). Behavioral recovery mirrored these structural improvements: nearly 60% of vehicle-treated DMD rats showed reduced activity and environmental disengagement (Grades 1–2), whereas both IROD cohorts maintained activity levels and grooming behaviors indistinguishable from WT controls (Supp. Fig. 1B). Porphyrin staining, a biomarker of chronic stress, was also reduced in a dose-dependent manner (Supp. Fig. 1C).

To complement these anatomical findings with functional assessments, we performed ex vivo contractile testing of the EDL. Untreated DMD rats exhibited a 2.75-fold deficit in absolute maximum tetanic force relative to WT controls, consistent with the extent of muscle wasting (Fig. 1F). Treatment with 3 mg/kg IROD restored absolute force 2.4-fold above untreated DMD controls, reaching levels statistically indistinguishable from WT. A parallel 2.4-fold improvement in absolute active stiffness was observed in the 3 mg/kg group relative to vehicle-treated DMD controls (Fig. 1G). However, when force and stiffness were normalized to muscle cross-sectional area (mCSA) (Supp. Fig. 1D), the functional differences between IROD-treated and untreated DMD groups were no longer significant (Fig. 1H–I). Specific tetanic force remained significantly lower in untreated DMD rats versus WT controls, but no significant difference was detected between the 3 mg/kg group and untreated DMD after normalization (Fig. 1H); normalized active stiffness showed the same pattern (Fig. 1I). These data indicate that IROD-mediated restoration of absolute contractile strength is driven primarily by an increase in total muscle mass rather than qualitative repair of individual myofibers, consistent with enhanced myofiber accretion as the dominant mechanism of functional rescue.

### Transcriptomic analysis reveals EP4 target engagement, sarcolemmal gene upregulation, and normalization of the myogenic program in IROD-treated muscle

To characterize the transcriptomic response to IROD treatment, we performed bulk RNA-seq analysis on skeletal muscle lysates harvested one week after the final dose. Principal component analysis (PCA) revealed clear separation between 3 mg/kg IROD-treated and vehicle-treated groups (Supp. Fig. 2A), and analysis of the top 40 differentially expressed genes further highlighted this distinctive transcriptional profile (Supp. Fig. 2B). Gene Ontology (GO) analysis demonstrated that the most significantly altered transcripts were associated with skeletal muscle contraction and developmental processes (Fig. 2A). The transcriptomic profile provided evidence consistent with sustained EP4-driven cAMP signaling within the dystrophic niche: significant upregulation of *Pde4a* and *Pde4b* was observed; as established negative feedback regulators of cAMP signaling, their upregulation is consistent with elevated intracellular cAMP activity (*18, 19*) (Fig. 2B). Significant downregulation of *Fbxo30* (*Musa1*), a key regulator of muscle atrophy (*20*), was also observed and is consistent with the muscle mass improvements shown in Fig. 2 (Fig. 2B).

**Fig. 2.**
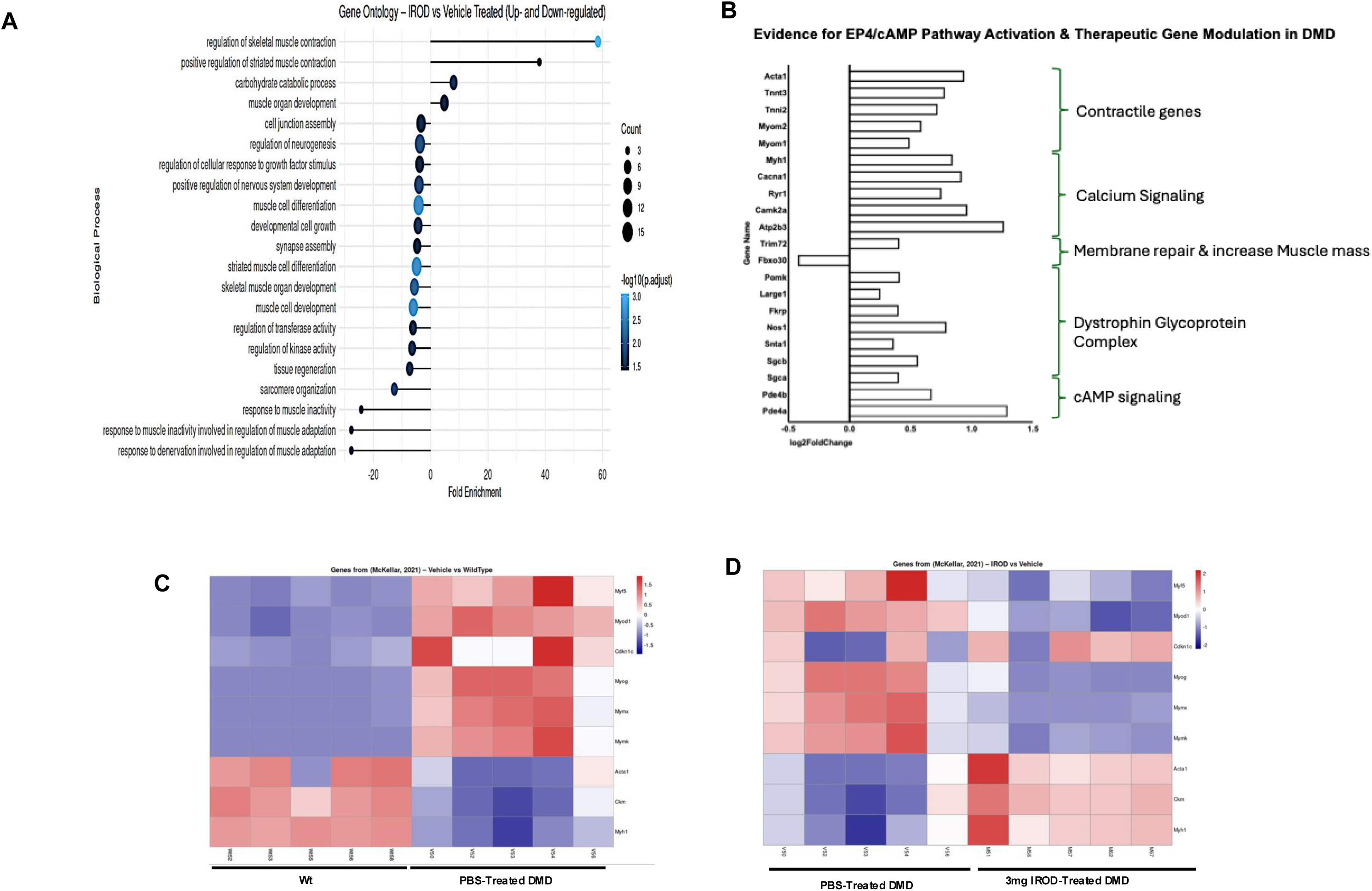
Transcriptomic analysis reveals EP4 target engagement, sarcolemmal gene upregulation, and normalization of the myogenic program in IROD-treated DMD muscle. (A) Gene Ontology (GO) biological process enrichment bubble plot from bulk RNA-seq of skeletal muscle lysates in 3 mg/kg IROD-treated versus vehicle-treated DMD rats. Bubble size represents the number of genes per term; color indicates −log₁₀(adjusted P-value). (B) Log₂ fold change of selected differentially expressed genes grouped by pathway and functional role, including cAMP signaling (*Pde4a*, *Pde4b*), muscle atrophy (*Fbxo30*), dystrophin-glycoprotein complex components (*Sgca*, *Sgcb*, *Snta1*, *Nos1*), sarcolemmal stabilization and membrane repair (*Trim72*, *Fkrp*, *Large1*, *Pomk*), calcium handling (*Atp2b3*, *Camk2a*, *Ryr1*, *Ryr2*, *Casq1*, *Cacna1s*), and the contractile apparatus (*Myh1*, *Mstn*, *Myom1*, *Myom2*, *Tnni2*, *Acta1*). (C) Heatmap of a nine-gene myogenic signature in age-matched WT versus untreated DMD rats, showing downregulation of mature structural markers (*Acta1*, *Ckm*, *Myh1*) and elevated myogenic progression markers in DMD muscle. (D) Heatmap of the same nine-gene signature comparing vehicle-treated and 3 mg/kg IROD-treated DMD rats. IROD reversed the pathological expression profile in (C), with downregulation of myogenic lineage genes (*Myod1*, *Myf5*, *Myf6*, *Myog*, *Mymk*, *Mym*) and a concomitant rise in mature structural markers. Color scale represents row-normalized Z-score. n = 5 per group; differentially expressed genes defined as |log₂FC| > 1, adjusted P < 0.05.

IROD treatment was associated with upregulation of multiple components of the dystrophin-glycoprotein complex (DGC), which is characteristically downregulated in DMD (*21*). Significant upregulation was observed for sarcoglycan subunits (*Sgca*, *Sgcb*), syntrophin (*Snta1*), and neuronal nitric oxide synthase (*Nos1*) (Fig. 2B). This was accompanied by upregulation of the membrane repair gene *Trim72* (Mitsugumin-53) (*22*) and DGC glycosylation genes—*Fkrp*, *Large1*, and *Pomk*—which are essential for sarcolemmal stabilization (*23*) (Fig. 2B). In addition, broad upregulation of genes involved in calcium handling (*Atp2b3*, *Camk2a*, *Ryr1*, *Ryr2*, *Casq1*, *Cacna1s*) (*24*) and the contractile apparatus (*Myh1*, *Mstn*, *Myom1*, *Myom2*, *Tnni2*, *Acta1*) (*25*) was observed, consistent with the restored contractile function documented in Fig. 1F–G (Fig. 2B).

To evaluate the state of the regenerative niche, we analyzed a signature set of nine genes involved in muscle stem cell proliferation and differentiation. Comparison of age-matched WT and untreated DMD rats revealed a pathological divergence in this gene set (Fig. 2C). Markers of mature muscle structure and function, including *Acta1*, *Ckm*, and *Myh1* were significantly downregulated in DMD muscle, while markers of active myogenic progression were concurrently elevated (*26, 27*), reflecting the chronic regenerative cycling characteristic of dystrophic tissue (Fig. 2C). Treatment with 3 mg/kg IROD reversed this pathological expression profile (Fig. 2D). Six genes involved in the myogenic lineage progression and myocyte fusion—*Myod1*, *Myf5*, *Myf6*, *Myog*, *Mymk*, and *Mym*—were significantly downregulated relative to untreated DMD controls, with a concomitant rise in mature structural markers (Fig. 2D). These findings are consistent with a reduction in chronic regenerative cycling and stabilization of the dystrophic muscle transcriptome toward a more mature phenotype.

### IROD treatment dose-dependently preserves skeletal muscle integrity and mitigates fibro-fatty remodeling in DMD rats

To determine whether the observed transcriptomic shifts and systemic improvements translated into structural preservation of skeletal muscle, we performed detailed histological analysis of the tibialis anterior (TA). The dose-dependent increase in TA muscle mass documented in Fig. 2C correlated with a marked rescue of myofiber morphology (Fig. 3A–D). Analysis of myofiber size distribution revealed that untreated DMD muscle showed a pathological shift toward smaller fibers, whereas IROD treatment partially rescued such shift in a dose-dependent manner, with a mode of 30–40 μm resembling the WT distribution (Fig. 3B). Quantification of average minimum Feret diameter confirmed that both the 1 mg/kg and 3 mg/kg cohorts exhibited significantly greater myofiber sizes compared with both untreated DMD controls and WT animals (Fig. 3C). IROD also dose-dependently reduced myofiber size heterogeneity, with the 3 mg/kg dose significantly lowering the coefficient of variation (CV) compared with the vehicle group, indicating a more uniform muscle architecture (Fig. 3D).

**Fig. 3.**
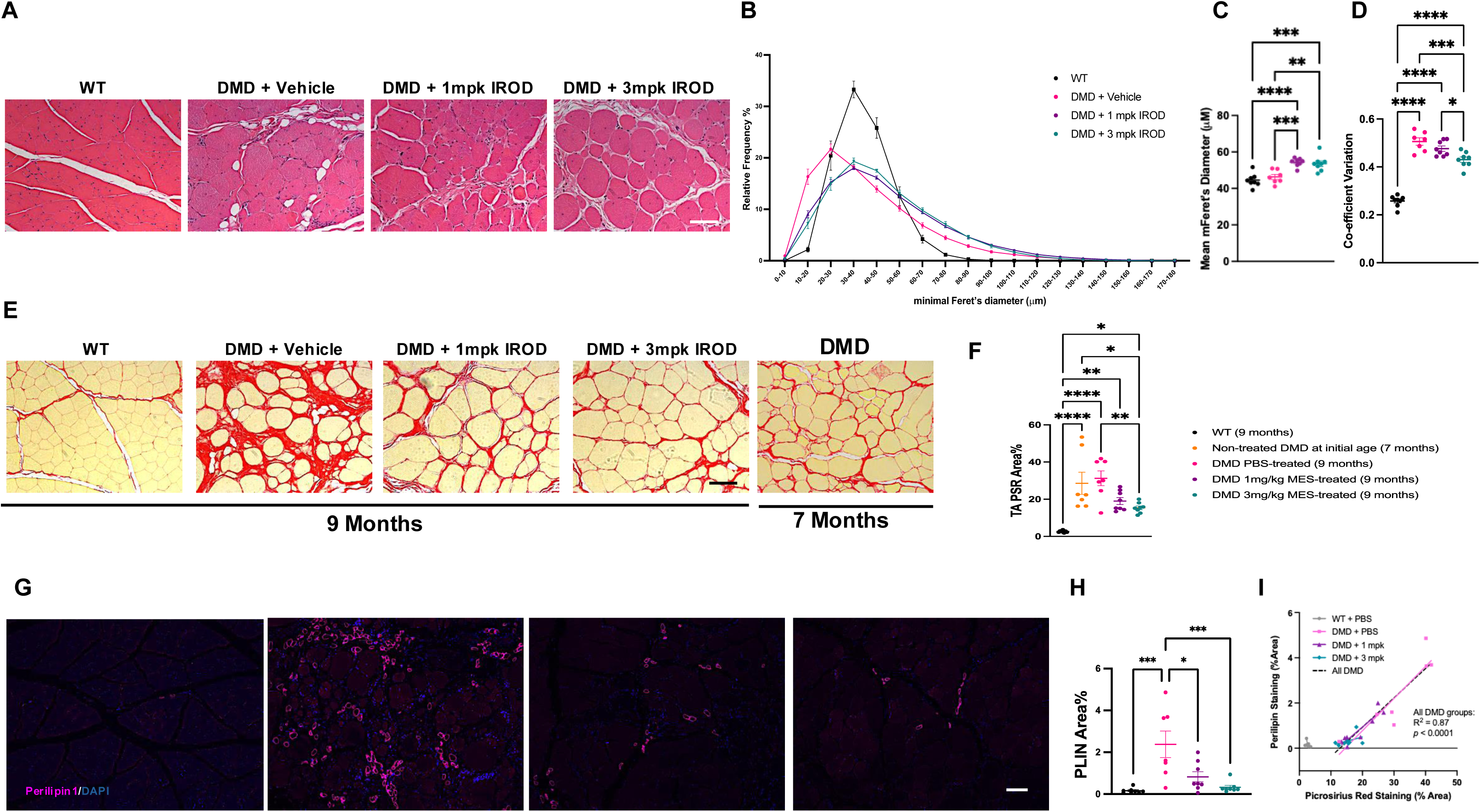
Dose-dependent rescue of myofiber morphology, reduction of fibrosis, and mitigation of intramuscular fat accumulation in the tibialis anterior of IROD-treated DMD rats. (A) Representative H&E-stained TA cross-sections from WT, 0 mg/kg, 1 mg/kg, and 3 mg/kg DMD groups. Scale bar = 100 μm. (B) Frequency distribution of myofiber minimum Feret diameter across groups. (C) Average myofiber minimum Feret diameter across groups. (D) Coefficient of variation (CV) of myofiber minimum Feret diameter as a measure of size heterogeneity. (E) Representative Picro-Sirius Red (PSR)-stained TA cross-sections. Scale bar = 100 μm. (F) PSR-positive area as a percentage of total TA cross-sectional area, including a pre-treatment baseline cohort of 7-month-old untreated DMD rats. Vehicle-treated DMD rats showed 31.2 ± 10.6% PSR-positive area; the 3 mg/kg group exhibited PSR levels significantly below the pre-treatment baseline (P < 0.05). (G) Representative perilipin-stained TA cross-sections. Scale bar = 100 μm. (H) Perilipin-positive area as a percentage of total TA cross-sectional area. Untreated DMD rats showed 2.37 ± 1.68% intramuscular fat accumulation; the 3 mg/kg group was reduced to levels statistically indistinguishable from WT controls. (I) Simple linear regression of PSR-positive fibrosis versus perilipin-positive intramuscular fat accumulation across all DMD rats (R² = 0.87, P < 0.0001). For dot plots (C, D, F, H), data are mean ± SEM; n = 5–6 per group. All statistical comparisons by two-way ANOVA; *P < 0.05, **P < 0.01, ***P < 0.001.

In addition to fiber morphology, we assessed the impact of IROD on the chronic fibrosis that characterizes DMD (Fig. 3E–F) (*28*). At 9 months of age, vehicle-treated DMD rats showed severe fibrosis, with Picro-Sirius Red (PSR) staining occupying 31.2 ± 10.6% of the TA muscle area (Fig. 3F). IROD dose-dependently reduced PSR-based fibrosis, a reduction that was consistent across the entire muscle belly (Supp. Fig. 4A). Notably, when compared with a baseline pre-treatment cohort of 7-month-old untreated DMD rats, the 3 mg/kg group exhibited PSR levels significantly below the pre-treatment starting point (Fig. 3F), consistent with active resolution of established fibrotic tissue at the higher dose.

This structural remodeling extended to intramuscular fat accumulation, assessed by perilipin staining (Fig. 3G–H). Untreated DMD rats displayed significantly elevated muscle fat (2.37 ± 1.68%), and IROD dose-dependently reduced intramuscular fat accumulation (Fig. 3H). In the 3 mg/kg group, adipocyte numbers were reduced to levels statistically indistinguishable from WT controls (Fig. 3H). A simple linear regression across all DMD rats revealed a strong positive correlation between fibrosis and intramuscular fat accumulation (R² = 0.87, *P* < 0.0001) (Fig. 3I), suggesting that IROD targets common pathological drivers or effectors of fibro-fatty muscle replacement.

Finally, we evaluated the impact of IROD on the cardiorespiratory system (Supp. Fig. 3B–D) (*29*). Histological analysis of the diaphragm revealed significant fibrosis across all DMD groups relative to WT (Supp. Fig. 3B), with no significant differences between treatment doses within this 8-week window. In the heart, all DMD groups showed significantly higher fibrosis than WT controls; however, treatment with 3 mg/kg IROD significantly reduced relative cardiomegaly—heart weight adjusted for body weight, compared with vehicle-treated DMD animals (Supp. Fig. 3C–D). These data indicate that while IROD mitigates pathological cardiac remodeling within 8 weeks, resolution of established fibrosis in cardiorespiratory tissues may require a longer treatment duration than that observed in hindlimb skeletal muscle.

### IROD intervention in young DMD rats modulates myofiber identity without rescuing muscle mass or architecture

To determine whether the fibrosis of the diaphragm that was resistant to late-stage treatment could be prevented by earlier intervention, we first characterized the natural history of fibrosis of the diaphragm in the DMD rat at 2, 4, 7, and 9 months of age compared with age-matched WT controls—a time course not previously reported in this model. Fibrosis of the diaphragm increased progressively and significantly at all time points relative to age-matched WT controls (Supp. Fig. 4A), confirming that pathological remodeling is present but less established at 2 months of age than in adolescent animals. We therefore initiated an early-intervention study in 2-month-old DMD rats, treating animals with weekly subcutaneous injections of vehicle (0 mg/kg) or 3 mg/kg IROD for 8 weeks alongside age-matched WT controls.

IROD treatment did not rescue any structural or functional endpoint in this cohort (Fig. 4B–I). In vivo grip strength assessment confirmed the absence of a functional effect of IROD and revealed that a significant deficit in muscle strength relative to WT controls becomes detectable by 4 months of age in this model (Supp. Fig. 4B). No significant drug-induced changes were observed in total body weight (Fig. 4B) or the absolute mass of the tibialis anterior (TA) (Fig. 4C), extensor digitorum longus (EDL) (Fig. 4D), or soleus muscles (Fig. 4E). Analysis of relative TA mass normalized to final body weight revealed a significantly higher TA/body weight ratio in DMD compared with WT controls (Supp. Fig. 4C), an observation that may be partly attributable to the significant reduction in fat mass, reflected by lower epididymal fat weight, in DMD rats relative to WT (Supp. Fig. 4D). Notably, 2-month-old DMD rats already exhibited a significant reduction in TA minimum Feret diameter and a significant increase in Picro-Sirius Red (PSR)-positive fibrosis compared with WT controls; IROD treatment did not rescue either deficit (Fig. 4F–G). Relative heart weight was not significantly different across groups at this stage (Fig. 4H), and fibrosis of the diaphragm was similarly unaffected by treatment (Fig. 4I).

**Fig. 4.**
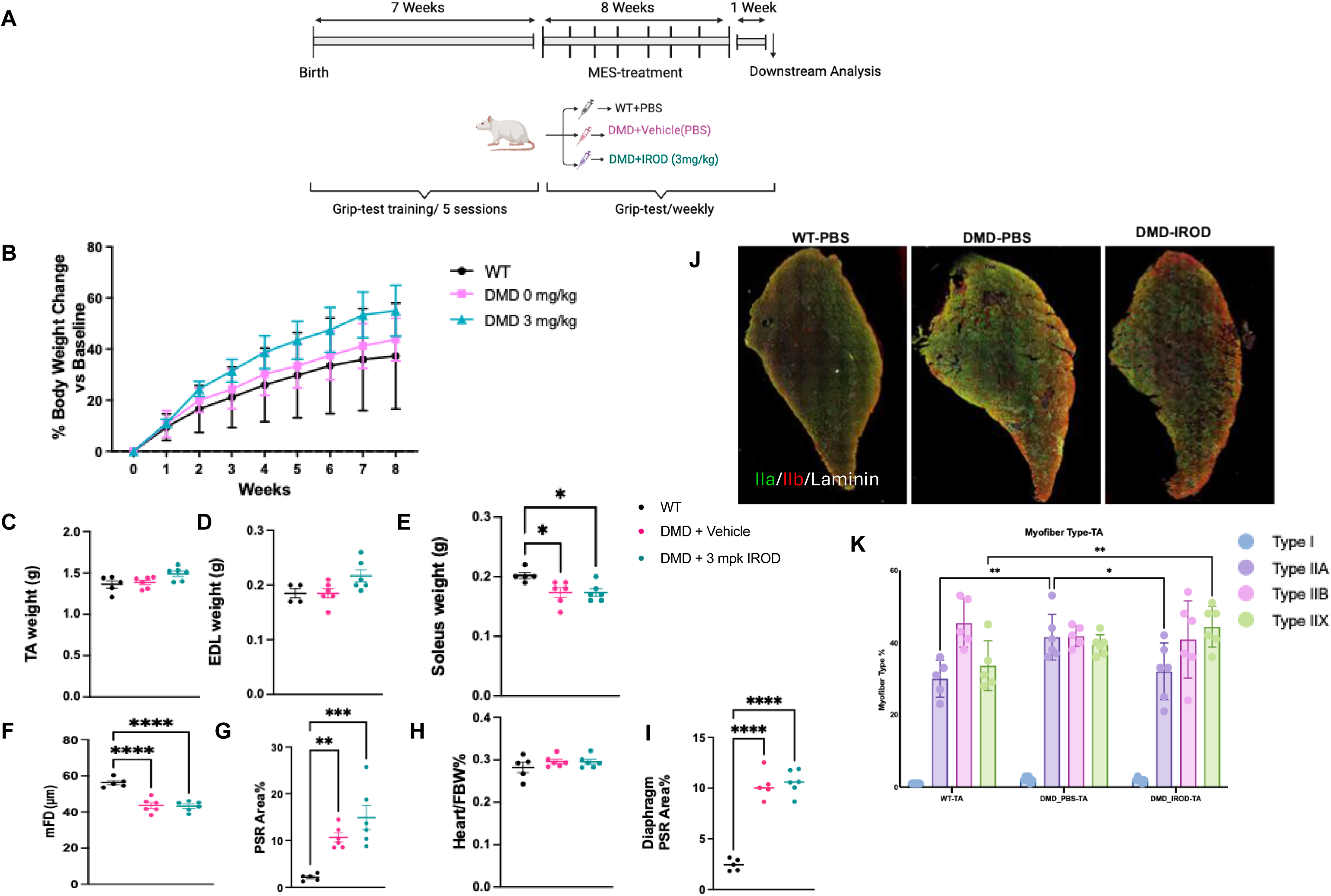
Early IROD intervention in 2-month-old DMD rats does not rescue muscle mass or architecture but significantly shifts TA myofiber type composition toward WT proportions. (A) Schematic of the early-intervention study design. Two-month-old DMD and WT rats received weekly subcutaneous injections of vehicle (0 mg/kg) or 3 mg/kg IROD for 8 weeks. (B) Percentage change in body weight from baseline. Data are mean ± SEM; n = 5–6 per group. (C–E) Absolute TA (C), EDL (D), and soleus (E) muscle weights at study end. No significant differences were observed between IROD-treated and vehicle-treated DMD groups. (F) TA myofiber minimum Feret diameter (mFD). IROD did not significantly rescue the reduction in fiber size in DMD rats. (G) PSR-positive area as a percentage of total TA cross-sectional area. IROD did not significantly reduce fibrosis relative to vehicle-treated DMD controls. (H) Relative heart weight (heart weight normalized to final body weight). No significant differences were observed across groups. (I) Diaphragm fibrosis quantified as PSR-positive area as a percentage of total diaphragm cross-sectional area. IROD did not significantly affect fibrosis of the diaphragm within the 8-week treatment window. (J) Representative immunostaining of whole TA muscle cross-sections illustrating myofiber type distribution across groups. (K) Myofiber type proportions in TA muscle. IROD treatment significantly restored the Type 2A fiber proportion to levels statistically indistinguishable from WT controls, with no significant changes in other fiber types. For dot plots (C–I), data are mean ± SEM; n = 5–6 per group. All statistical comparisons by two-way ANOVA; *P < 0.05, **P < 0.01, ***P < 0.001.

Despite the absence of gross structural rescue, we investigated whether IROD exerted cellular-level affects myofiber identity. In DMD patients, Type II fast-twitch fibers are preferentially susceptible to mechanical stress-driven degeneration, a selective vulnerability that precedes global muscle atrophy (*30*). As myofiber type composition in the DMD rat has not previously been characterized, we assessed fiber type distribution in three limb muscles and evaluated the effect of IROD treatment on myofiber identity. No significant fiber-type changes were observed in the EDL or soleus (Supp. Fig. 4E–F). In the TA, however, immunostaining of full muscle cross-sections revealed a clear reorganization of myofiber type composition in IROD-treated DMD rats toward the WT pattern, in contrast to the pathological fiber type distribution of untreated DMD animals (Fig. 4J). Quantification confirmed a significant restoration of Type 2A fiber relative abundance to levels statistically indistinguishable from WT controls in IROD-treated rats (Fig. 4K). These findings demonstrate that IROD exerts a meaningful effect on myofiber identity at this early disease stage, remodeling the cellular composition of dystrophic muscle toward a healthier phenotype even in the absence of gross structural rescue.

### IROD drives myofiber pool restoration and inhibits fibro-adipogenic progenitor differentiation in adolescent DMD muscle

To further characterize the cellular drivers underlying the stage-specific efficacy of IROD in adolescent DMD rats, we performed an in-depth histological analysis of TA muscles from 9-month-old treated animals. The total number of myofibers in the TA of untreated DMD rats was 45.6% lower than that of WT controls (Fig. 5A). IROD treatment significantly increased total myofiber count in a dose-dependent manner, and the 3 mg/kg group achieved a mean total myofiber number statistically indistinguishable from WT controls, indicating near-complete rescue of the fiber deficit (Fig. 5A). To determine whether this increase was driven by active regeneration, we quantified centronucleated myofibers, an established histological marker of newly fused muscle cells (*31*) (Fig. 5B). While WT muscles showed negligible centronucleation (∼0.5%), untreated DMD rats exhibited significantly higher proportions of centrally located nuclei (21.5 ± 3.9%), reflecting chronic compensatory regeneration. Treatment with 3 mg/kg IROD further and significantly increased the percentage of centronucleated myofibers compared with vehicle-treated DMD controls (Fig. 5B). This regenerative response was corroborated by a significant increase in embryonic myosin heavy chain (eMHC)-positive myofibers in treated groups (Fig. 5C–D). These eMHC^+^ fibers appeared in distinct clustered patterns, consistent with synchronized, coordinated regeneration. While this clustered pattern is characteristic of early-stage disease in 4-month-old DMD rats (Supp. Fig. 5A), it was largely absent in untreated 9-month-old animals where regenerative capacity is typically diminished; treatment with IROD restored eMHC+ fibres in 9-month-old DMD muscle (Supp. Fig. 5A). These observations are consistent with an age-dependent effect on regenerative capacity. At 2–4 months of age, eMHC+ fibres are already elevated (Supp. Fig. 5A), leaving limited capacity for further drug-induced increases (Supp. Fig. 5B). At 7–9 months, the data are consistent with IROD re-activating a regenerative program that is otherwise suppressed in chronically wasting adolescent muscle, enabling restoration of the total myofiber pool to WT levels (Fig. 5A–B).

**Fig. 5.**
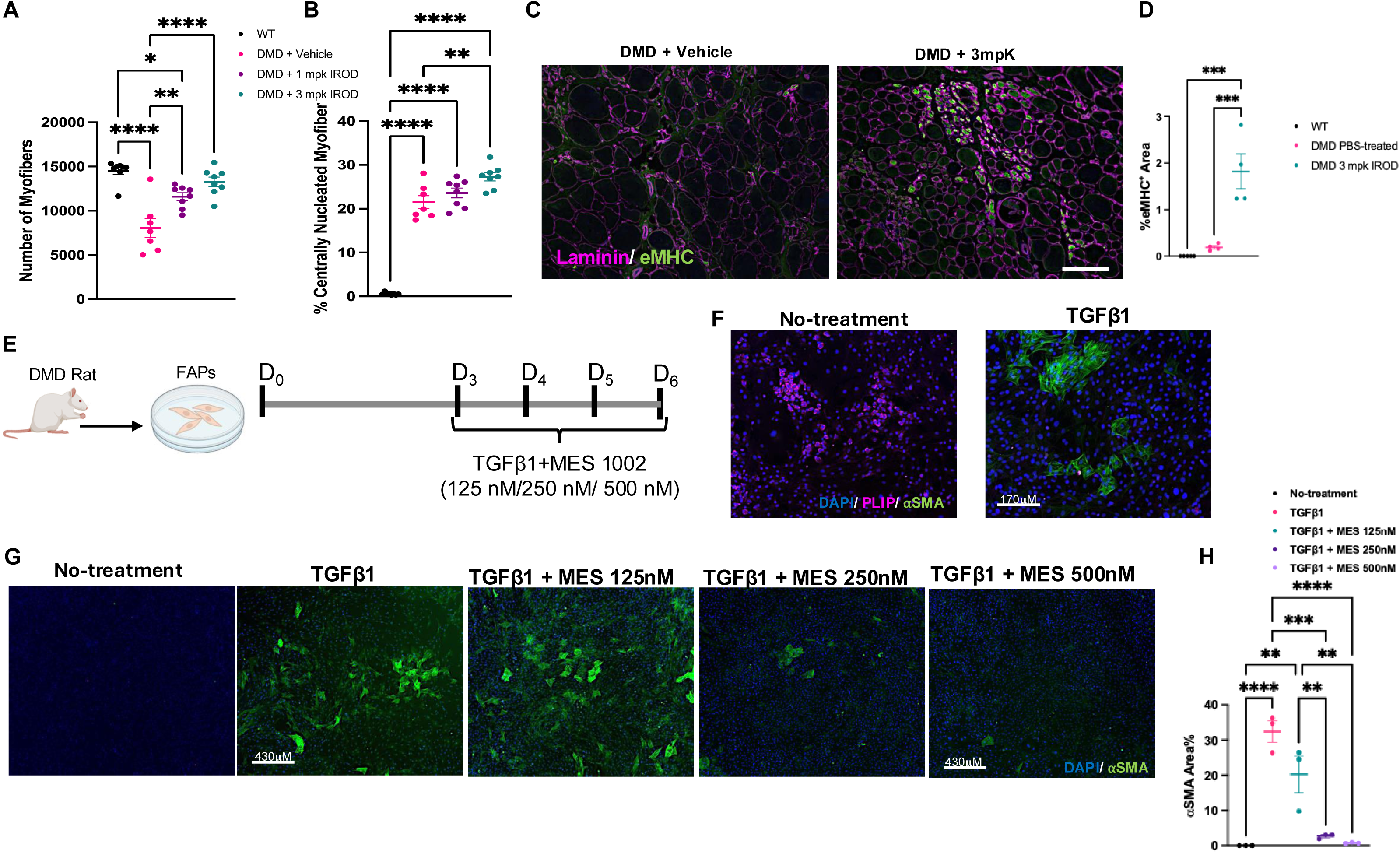
IROD drives myofiber pool restoration and inhibits fibro-adipogenic progenitor differentiation in adolescent DMD muscle. (A) Total myofiber count in the TA of WT, 0 mg/kg, 1 mg/kg, and 3 mg/kg DMD groups at 9 months. Untreated DMD rats exhibited a 45.6% reduction relative to WT; the 3 mg/kg group reached a count statistically indistinguishable from WT. (B) Percentage of centronucleated myofibers across groups. WT muscles showed negligible centronucleation (∼0.5%); untreated DMD rats exhibited 21.5 ± 3.9%. Treatment with 3 mg/kg IROD further and significantly increased centronucleation relative to vehicle-treated controls (P = 0.0023). (C) Representative immunofluorescence images of eMHC-stained TA cross-sections. eMHC+ fibers appear in clustered patterns in IROD-treated animals, consistent with synchronized regeneration. Scale bar = 50 μm. (D) Percentage of eMHC-positive myofibers across groups. For dot plots (A, B, D), data are mean ± SEM; n = 7–8 per group. (E) Schematic of the in vitro FAP isolation and differentiation assay. FAPs were isolated from 7-month-old DMD rats and cultured under adipogenic or TGF-β-induced fibrogenic conditions, with or without co-treatment with MES1002. (F) Representative immunofluorescence images of FAP differentiation under adipogenic and TGF-β-induced fibrogenic conditions. Scale bar = 170 μm. (G) Representative α-SMA immunofluorescence in TGF-β-stimulated FAPs co-treated with vehicle or MES1002 at 125, 250, and 500 μM. Scale bar = 430 μm. (H) Quantification of α-SMA-positive area as a percentage of total cell area. MES1002 dose-dependently inhibited FAP differentiation into α-SMA-positive myofibroblasts. Data are mean ± SEM; n = 3 independent experiments. All statistical comparisons by two-way ANOVA; *P < 0.05, **P < 0.01, ***P < 0.001.

This drug-induced regeneration was not accompanied by exacerbation of inflammation or sarcolemmal injury. Staining for CD68, a marker of macrophages and inflammatory cells, revealed minimal-to-moderate presence across all DMD groups with no significant differences between IROD-treated and vehicle-treated rats (Supp. Fig. 5C–D). Sarcolemmal integrity was assessed by cytoplasmic IgG staining, which identifies myofibers undergoing active membrane disruption (Supp. Fig. 5E–F). Approximately 1% of TA myofibers were IgG-positive across all DMD groups, and IROD treatment did not significantly alter this proportion relative to untreated DMD controls. No IgG-positive fibers were detected in WT rats (Supp. Fig. 5F). These findings indicate that IROD promotes myofiber pool expansion without aggravating inflammation or compromising sarcolemmal integrity. In addition, they support the notion that IROD acts by enhancing regeneration, but does not prevent the myofibre damage typical of DMD.

To investigate the cellular mechanism underlying the observed resolution of fibrosis, we evaluated the direct impact of MES-1002, the active EP4 agonist moiety released upon hydrolysis of the IROD prodrug, on fibro-adipogenic progenitors (FAPs) isolated from 7-month-old DMD rats (Fig. 5E–H). Unlike IROD, which requires esterase-mediated hydrolysis of the linker to release the active EP4 agonist moiety, MES1002 can be applied directly in vitro to assess cell-specific EP4 responses independently of the prodrug targeting mechanism. In the absence of exogenous stimulation, FAPs isolated from DMD muscle primarily exhibited an adipogenic differentiation program; however, treatment with TGF-β triggered a significant shift toward a fibrogenic phenotype characterized by α-SMA expression and collagen production (Fig. 5F). FAPs co-treated with MES-1002 at 125, 250, and 500 μM showed dose-dependent inhibition of differentiation into α-SMA-positive myofibroblasts (Fig. 5G–H). These data demonstrate for the first time that EP4 agonism directly inhibits the fibrogenic differentiation of FAPs derived from dystrophic muscle, providing a cellular mechanism for the anti-fibrotic effects of IROD observed in vivo.

### IROD selectively targets dystrophic muscle

Given that PGE2 signaling through the EP4 receptor is essential for muscle satellite cell expansion and regeneration (10), we investigated the mechanism by which IROD exerts its effects on dystrophic skeletal muscle. Two non-mutually exclusive mechanisms were considered: release of the active EP4 agonist MES1002 from bone depots with subsequent permeation into adjacent muscle tissue, and direct accumulation of the intact prodrug in damaged muscle driven by either increased sarcolemmal permeability or binding to the micro-calcifications that accumulate in dystrophic myofibers as a consequence of chronic membrane disruption and calcium dyshomeostasis (31).

To distinguish between these possibilities, we assessed IROD biodistribution using two complementary radiotracer approaches. Subcutaneous administration of tritium-labeled IROD, with the tritium label placed on the EP4 agonist moiety MES1002, revealed slightly increased accumulation in kidney and liver of DMD rats relative to WT controls, while bone distribution was comparable between genotypes (Fig. 6A). In hindlimb skeletal muscle, DMD rats exhibited markedly elevated blood-normalized tissue concentrations relative to WT controls across all examined muscle groups, with no correlation between drug concentration and anatomical proximity to bone (Fig. 6B), consistent with vascular delivery as the dominant distribution mechanism rather than passive diffusion from adjacent skeletal tissue.

**Fig. 6.**
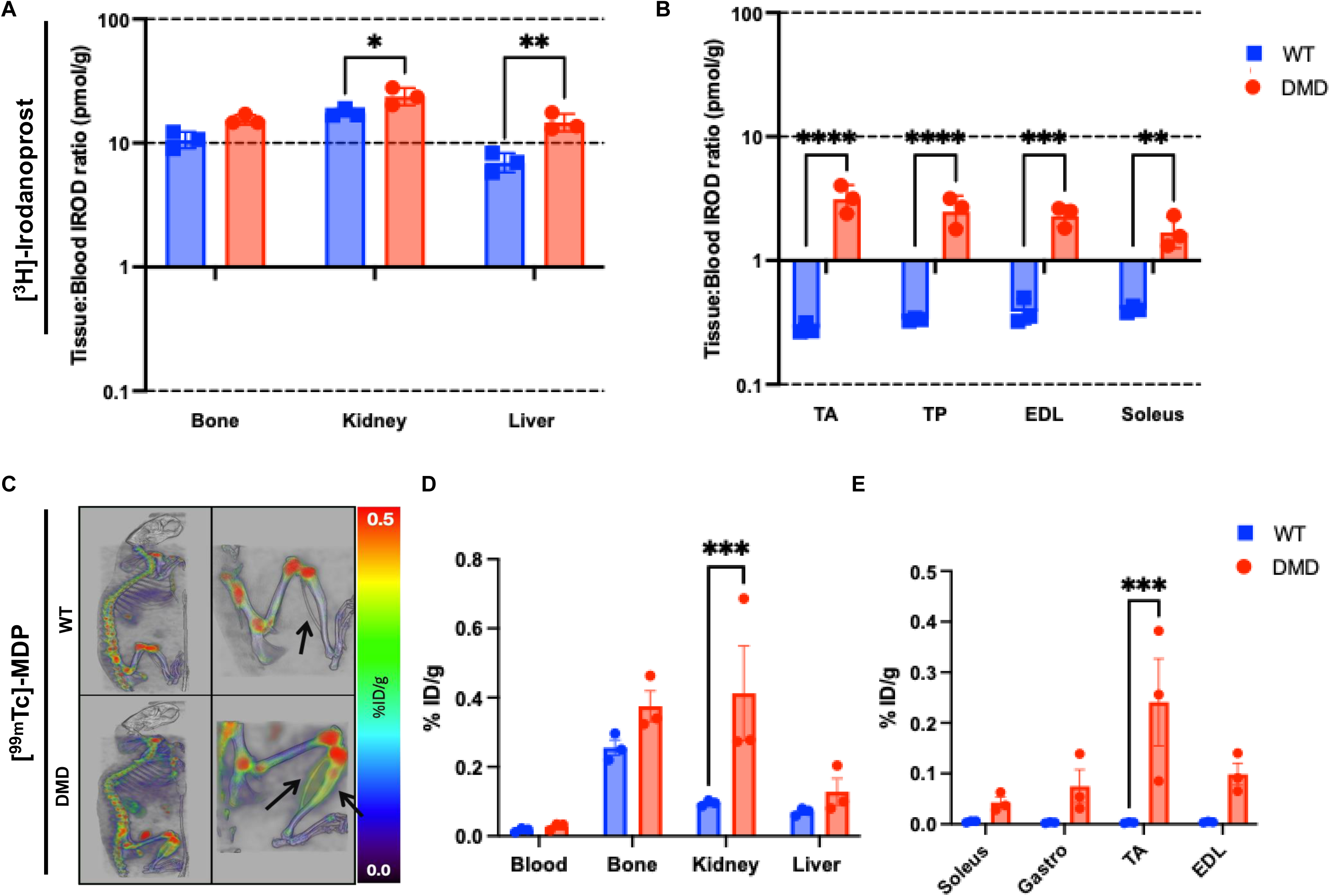
IROD Tissue distribution of [³H]-Irodanoprost and ⁹⁹ᵐTc-MDP reveals preferential accumulation in dystrophic muscle. **(A)** Tissue:Blood [³H]-Irodanoprost ratio (pmol/g) in bone, kidney, and liver of WT and DMD rats at 24 h after subcutaneous administration. Data are mean ± SEM; n = 3 per group. Y-axis is displayed on a log scale. **(B)** Tissue:Blood [³H]-Irodanoprost ratio (pmol/g) in individual hindlimb muscles of WT and DMD rats. The absence of a significant correlation between drug concentration and anatomical proximity of limb muscles to bone is consistent with vascular delivery as the dominant distribution mechanism. Y-axis is displayed on a log scale. Data are mean ± SEM; n = 3 per group. Y-axis is displayed on a log scale. **(C)** Representative SPECT/CT images of WT and DMD rats at 5 h post-injection. Right panels show magnified views of the hindlimb region. Images are displayed on a common %ID/g scale. **(D)** Quantification of ⁹⁹ᵐTc-MDP uptake (%ID/g) in blood, bone, kidney, and liver of WT and DMD rats from ex vivo tissue counting following SPECT/CT imaging. **(E)** Quantification of ⁹⁹ᵐTc-MDP uptake (%ID) in individual hindlimb muscles of WT and DMD rats. Data are mean ± SEM; n = 3 per group. Data are mean ± SEM; n = 3 per group. TA: tibialis anterior; TP: tibialis posterior; EDL: extensor digitorum longus; Gastro: gastrocnemius. All statistical comparisons by two-way ANOVA with post-hoc Tukey test; *P < 0.05, ***P < 0.001, ****P < 0.0001.

Because the tritium label was placed on the EP4 agonist moiety of IROD, which is released upon hydrolysis of the ester linker, these data could not distinguish between muscle accumulation of intact [³H]-IROD and free MES1002. We therefore independently evaluated the affinity of the phosphonate moiety for dystrophic muscle using ⁹⁹ᵐTc-MDP (Fig. 6B), a clinically established phosphonate radiotracer with well-characterized hydroxyapatite affinity (*32*). While WT animals showed the expected bone-dominated distribution with minimal muscle signal, DMD rats exhibited pronounced hindlimb tracer accumulation visible on SPECT/CT imaging (Fig. 6C) and confirmed by ex vivo quantification of individual muscles (Fig. 6D–E). Together, these data establish that IROD can efficiently reach damaged muscle, providing a mechanistic basis for the systemic musculoskeletal rescue observed with treatment.

## Discussion

A critical limitation in the current landscape of DMD therapeutics is the narrow eligibility window for emerging interventions. Although the recent FDA approval of delandistrogene moxeparvovec (Elevidys) expanded gene therapy access to patients aged 4 years and older, this approval has subsequently been restricted to ambulatory patients following reports of fatal acute liver failure in non-ambulatory individuals, further narrowing the eligible population (*33*).

Moreover, even within the ambulatory population, the clinical benefit of micro-dystrophin gene therapy has not been consistently demonstrated against placebo in phase 3 trials (*34*). Adolescent patients, who often present with advanced fibro-fatty replacement, established fibrosis, and depleted regenerative capacity, have substantially fewer therapeutic options. Current standard-of-care corticosteroids partially mitigate inflammatory burden but do not reverse established fibrosis, restore myofiber number, or reactivate a stalled myogenic program, and are associated with significant adverse effects including osteoporosis and fragility fractures with long-term use (*2, 9*). Our study directly addresses this unmet need by utilizing the DMD rat model, which more accurately recapitulates the severe adolescent human phenotype than the standard mdx mouse (*16*). The data presented here demonstrate that the therapeutic window for IROD is well-suited to this advanced disease stage: in 7-month-old adolescent animals, we observed a dose-dependent rescue of systemic parameters and muscle architecture that was not replicated in younger, 2-month-old cohorts.

The biodistribution data provide insight into how a bone-targeted prodrug achieves meaningful and selective muscle exposure. Although the bisphosphonate moiety was designed for skeletal mineral affinity, biodistribution studies revealed elevated uptake across all examined skeletal muscles in DMD rats relative to WT controls, with no correlation between uptake and proximity to bone across individual hindlimb muscles. This pattern is consistent with vascular delivery and enhanced permeability of diseased tissues as the dominant distribution mechanism but also with selective tissue retention mediated by binding to the micro-calcifications that are known to accumulate within dystrophic myofibers as a consequence of chronic membrane disruption and calcium dyshomeostasis (*35–37*).By exploiting these mineral deposits as disease-specific tissue anchors, IROD could establish a localized reservoir of active EP4 agonist within the most pathologically affected regions of muscle. This hypothetical mechanism implies that IROD efficacy may be self-amplifying: greater dystrophic burden, and therefore greater calcification, would be expected to drive greater drug accumulation and more sustained EP4 signaling at the sites of most severe damage. Proof of a role for calcifications in determining IROD biodistribution will require more definitive studies (*38*).

The observation that IROD modulated myofiber type composition in young animals, restoring the pathological reduction in Type 2A fiber proportion toward WT levels in the TA, provides two important insights. First, it confirms that the drug reaches and engages skeletal muscle at 2 months of age. Second, it provides a characterization of myofiber type composition in the DMD rat, demonstrating that the model recapitulates the preferential loss of Type II fast-twitch fibers documented in human DMD muscle biopsies (*30*), and that EP4 agonism can influence myofiber identity independently of gross structural remodeling. The absence of broader structural rescue in young animals is more readily explained by the regenerative state of the tissue at this age. At 2–4 months, endogenous regenerative activity is already near maximal, as evidenced by high baseline levels of eMHC+ fibers and clustered regeneration. EP4 signaling via the cAMP/phospho-CREB/Nurr1 axis is a well-established driver of satellite cell expansion and muscle regeneration (*11*), and under conditions of already-active endogenous regeneration, additional EP4-mediated pro-regenerative signaling may have limited capacity to further amplify a program that is already fully engaged. This interpretation reframes the stage-dependent response not as a failure of drug delivery in young animals, but as a ceiling effect imposed by the regenerative state of the tissue at the time of treatment.

A prevailing paradigm in DMD research holds that chronic degeneration eventually leads to depletion or dysfunction of the satellite cell pool in advanced disease (*39–41*). Our data in adolescent animals are consistent with an alternative interpretation: that the regenerative machinery may not be irreversibly exhausted, but rather is inhibited by the pathological microenvironment, and that such inhibition can be overcome by specific EP4-mediated cues. This is supported by recent evidence that aged muscle stem cells harbor blunted PGE2–EP4 signaling causing precocious commitment, and that PGE2 treatment reverses this dysfunction by altering chromatin accessibility and triggering cell cycle re-entry (*42*). In untreated 9-month-old animals, synchronized clustered regeneration was largely absent, consistent with suppression of the regenerative program in chronically wasting muscle. Following IROD treatment, clusters of eMHC+ fibres were restored, total myofiber count returned to levels statistically indistinguishable from WT controls, and centronucleated fiber proportions increased significantly. These observations are consistent with IROD re-engaging a latent regenerative program that is suppressed but not extinguished in adolescent DMD muscle.

The reduction of fibrosis below pre-treatment baseline levels in adolescent animals indicates that IROD facilitates active resolution of established fibrotic tissue, not merely prevention of further accumulation. FAPs are the primary source of collagen deposition in dystrophic muscle and are pathologically dysregulated in DMD, with elevated TGF-β signaling driving their differentiation into α-SMA-positive myofibroblasts (*43*). Targeting FAP fibrogenesis has emerged as a validated therapeutic strategy; the HDAC inhibitor givinostat, which restricts FAP-driven fibrosis in dystrophic muscle, has received regulatory approval for DMD (*44*). The in vitro data presented here extend this framework by demonstrating, for the first time, that EP4 agonism directly inhibits the fibrogenic differentiation of FAPs isolated from dystrophic muscle in a dose-dependent manner. When FAPs from 7-month-old DMD rats were challenged with TGF-β and co-treated with MES1002, the active moiety of IROD, differentiation into α-SMA-positive myofibroblasts was dose-dependently inhibited. This identifies EP4 signaling as a cell-autonomous regulator of FAP fate in the dystrophic context and provides a mechanistic basis for the fibrosis resolution observed in vivo. The convergence of FAP inhibition and satellite cell reactivation through a shared EP4/cAMP axis suggests that IROD simultaneously targets two important pathological drivers of the fibro-fatty niche: excessive collagen deposition and impaired myofiber regeneration. This pro-regenerative response was not accompanied by increased inflammation or sarcolemmal disruption, indicating that IROD promotes myofiber pool expansion without exacerbating tissue stress.

A strong linear correlation (R² = 0.87) between reduction of intramuscular fat accumulation and fibrosis resolution was identified across all DMD animals. This relationship is mechanistically consistent with the shared FAP origin of both pathological processes: FAPs are the cellular source of both myofibroblasts and intramuscular adipocytes in dystrophic muscle (*45*), and EP4-mediated inhibition of FAP differentiation would be expected to reduce both fibrogenic and adipogenic outputs simultaneously. This finding may have clinical relevance for patient monitoring. MRI-based fat fraction imaging has been validated as an objective biomarker of disease progression in DMD that correlates with functional outcomes and predicts clinically meaningful endpoints such as loss of ambulation independently of age (*46, 47*), and there is growing support for its use as a surrogate endpoint in DMD clinical trials (*48*). While direct assessment of muscle fibrosis requires invasive biopsies that are often impractical in advanced DMD, intramuscular fat infiltration can be non-invasively quantified using proton density fat fraction (PDFF) or Dixon MRI sequences. If the linear relationship between these two pathological components is confirmed in human DMD tissue, reduction in intramuscular fat could serve as a non-invasive surrogate biomarker for fibrosis resolution, providing a clinically accessible means of assessing anti-fibrotic efficacy without repeated histological intervention.

The physiological rescue is mirrored by a fundamental shift in the myogenic transcriptional landscape. Bulk RNA-seq analysis revealed significant downregulation of six genes central to the myogenic lineage and myocyte fusion, *Myod1*, *Myf5*, *Myf6*, *Myog*, *Mymk*, and *Mym*, which are chronically upregulated in DMD as muscle engages in sustained but ultimately ineffective degeneration-regeneration cycling. Their downregulation, accompanied by a concomitant rise in mature structural markers, is consistent with stabilization of the dystrophic transcriptome toward a more mature phenotype. The functional signature of IROD, restoration of absolute contractile force while specific force after normalization to cross-sectional area remains unchanged, underscores its primary mechanism as a pro-regenerative agent that expands the total myofiber pool rather than improving the resistance of individual fibres to damage or their intrinsic contractile quality. This distinction is particularly relevant for adolescent patients, in whom critical loss of muscle volume is the dominant driver of functional decline. Systemic improvements in behavioral activity and reduction of porphyrin-based stress markers further indicate that the structural rescue translates into a measurable improvement in overall wellbeing. At the cardiac level, treatment with 3 mg/kg IROD significantly reduced relative cardiomegaly despite the absence of significant cardiac fibrosis resolution within the 8-week treatment window, suggesting that EP4-mediated signaling may attenuate pathological cardiac remodeling independently of, and potentially preceding, fibrosis resolution. As IROD addresses the fibro-fatty niche and restores myofiber mass without directly targeting the underlying dystrophin deficit, it may be well-suited to combination with mutation-specific therapies such as exon-skipping oligonucleotides or micro-dystrophin gene replacement, where a more permissive structural environment could enhance the benefit of dystrophin restoration.

Several limitations of this study warrant consideration. Biodistribution and early-intervention cohorts included small sample sizes (n = 3 per genotype for radiotracer studies; n = 6 per group for the 2-month cohort), which limits statistical power in these specific analyses. While ^99m^Tc-MDP and [^3^H]-IROD distribution to limb muscles was similar we have not strictly demonstrated a significant correlation of microcalcification deposits with efficacy (*38*). All animals were male, consistent with the predominant sex affected by DMD, but sex-specific effects of IROD cannot be excluded. The DMD rat, while a strong model of the adolescent human phenotype, does not fully recapitulate all aspects of human DMD, and the translational relevance of the observed regenerative and anti-fibrotic responses will require validation in human tissue and ultimately in clinical studies. The in vitro FAP experiments demonstrate a direct cellular effect of MES1002 on fibrogenic differentiation but do not establish whether this mechanism accounts for the full extent of fibrosis resolution observed in vivo, where additional cell types and signaling pathways are likely to contribute. Fibrosis of the diaphragm and cardiac fibrosis were not significantly resolved within the 8-week treatment window, suggesting that longer treatment durations or higher doses may be required to address established fibrosis in cardiorespiratory tissues. Whether the progressive fibrosis of the diaphragm can be prevented or attenuated by prolonged intervention remains an important open question for future studies.

---

**Supplementary Fig. 1.**
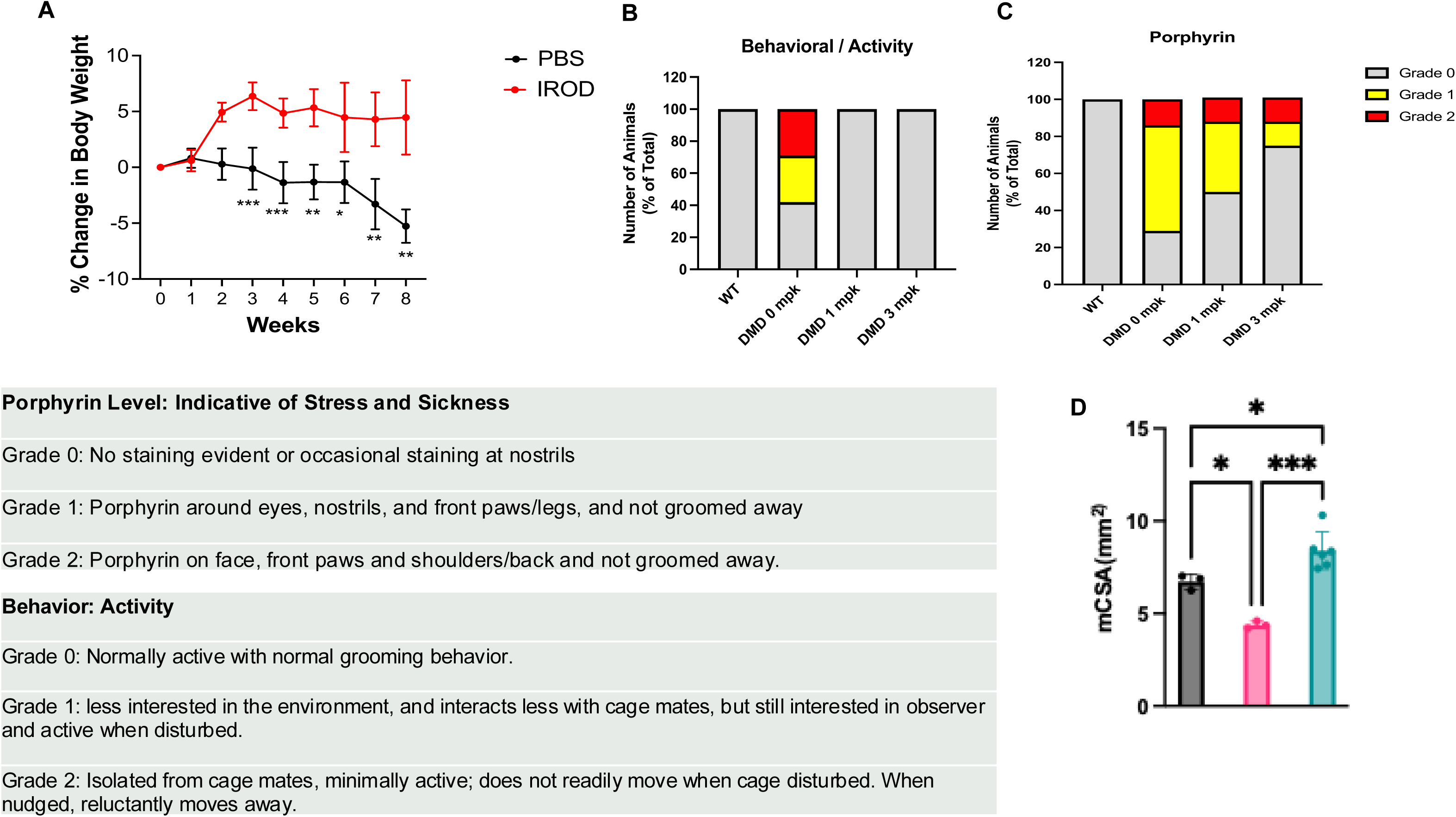
Behavioral health, porphyrin staining, and EDL muscle cross-sectional area in IROD-treated DMD rats. (A) Percentage change in body weight from baseline over the treatment period in DMD vehicle-treated (0 mg/kg) and DMD 3 mg/kg IROD-treated rats. Data are mean ± SD; n = 6–9 per group. (B) Behavioral health scores in WT, 0 mg/kg, 1 mg/kg, and 3 mg/kg DMD groups, graded from Grade 0 (normal activity) to Grade 2 (isolated, minimally active). Nearly 60% of vehicle-treated DMD rats scored Grade 1 or 2; both IROD-treated cohorts maintained activity indistinguishable from WT. Data are mean ± SEM; n = 7–8 per group. (C) Porphyrin staining grades in WT, 0 mg/kg, 1 mg/kg, and 3 mg/kg DMD groups, scored from Grade 0 (no staining) to Grade 2 (extensive staining). High-grade staining was present in 71% of vehicle-treated DMD rats, compared with 51% and 26% in the 1 mg/kg and 3 mg/kg groups, respectively. Data are mean ± SEM; n = 7–8 per group. (D) EDL muscle cross-sectional area (CSA) estimated as: CSA (mm²) = W (mg) / [D (mg/mm³) × L (mm)], where W is muscle wet weight, L is muscle length, and D is mammalian skeletal muscle density (1.06 mg/mm³). Data are mean ± SEM; n = 3–6 per group. All statistical comparisons by two-way ANOVA; *P < 0.05, **P < 0.01, ***P < 0.001.

**Supplementary Fig. 2.**
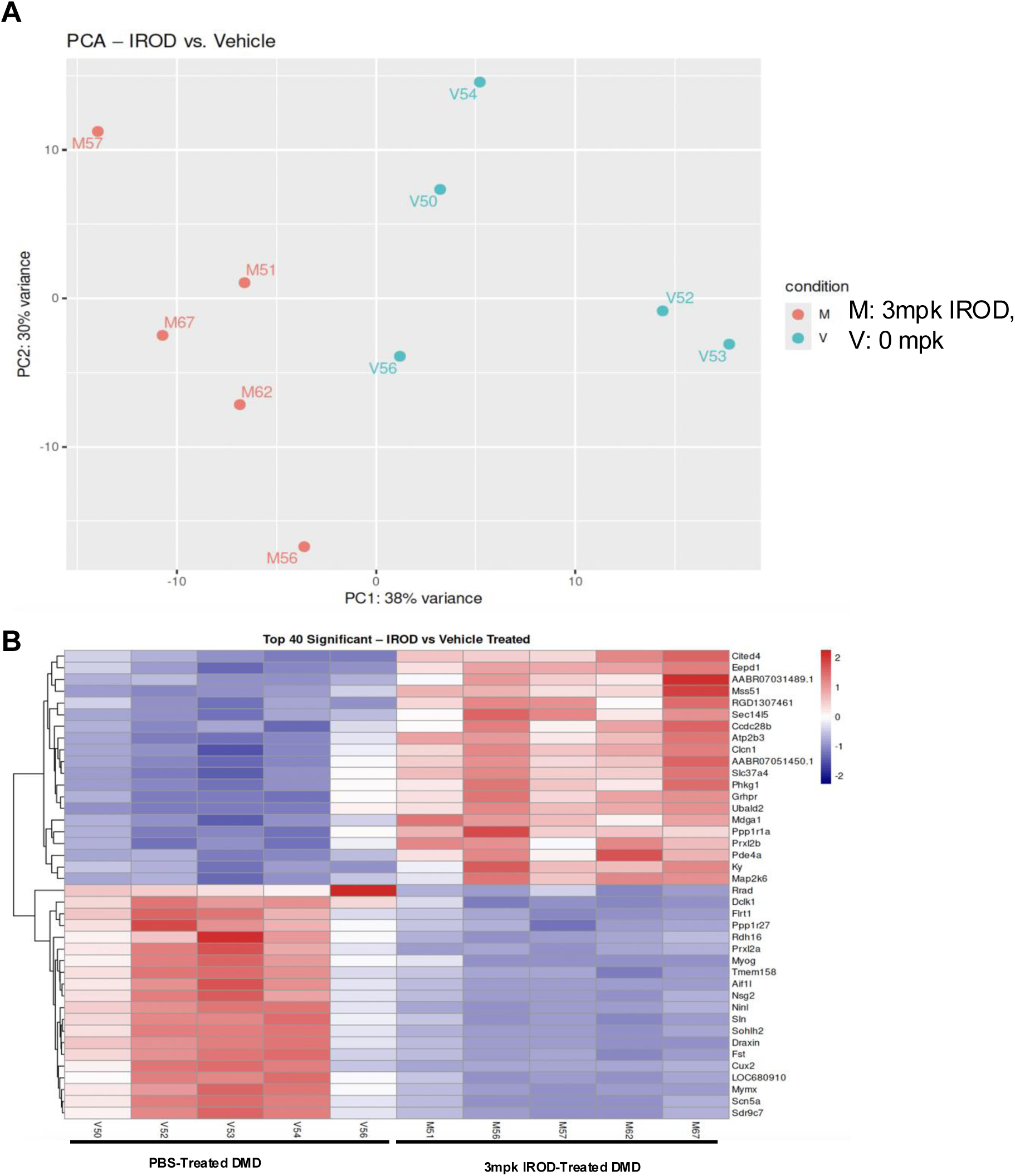
Global transcriptional separation and top differentially expressed genes in vehicle-treated versus 3 mg/kg IROD-treated DMD rats. (A) Principal component analysis (PCA) of bulk RNA-seq data from skeletal muscle lysates of vehicle-treated (0 mg/kg) and 3 mg/kg IROD-treated DMD rats harvested one week after the final dose. PC1 and PC2 account for 38% and 30% of total variance, respectively. n = 5 per group. (B) Heatmap of the top 40 differentially expressed genes between vehicle-treated and 3 mg/kg IROD-treated DMD rats, ranked by adjusted P-value. Color scale represents row-normalized Z-score. n = 5 per group; differentially expressed genes defined as |log₂FC| > 1, adjusted P < 0.05.

**Supplementary Fig. 3.**
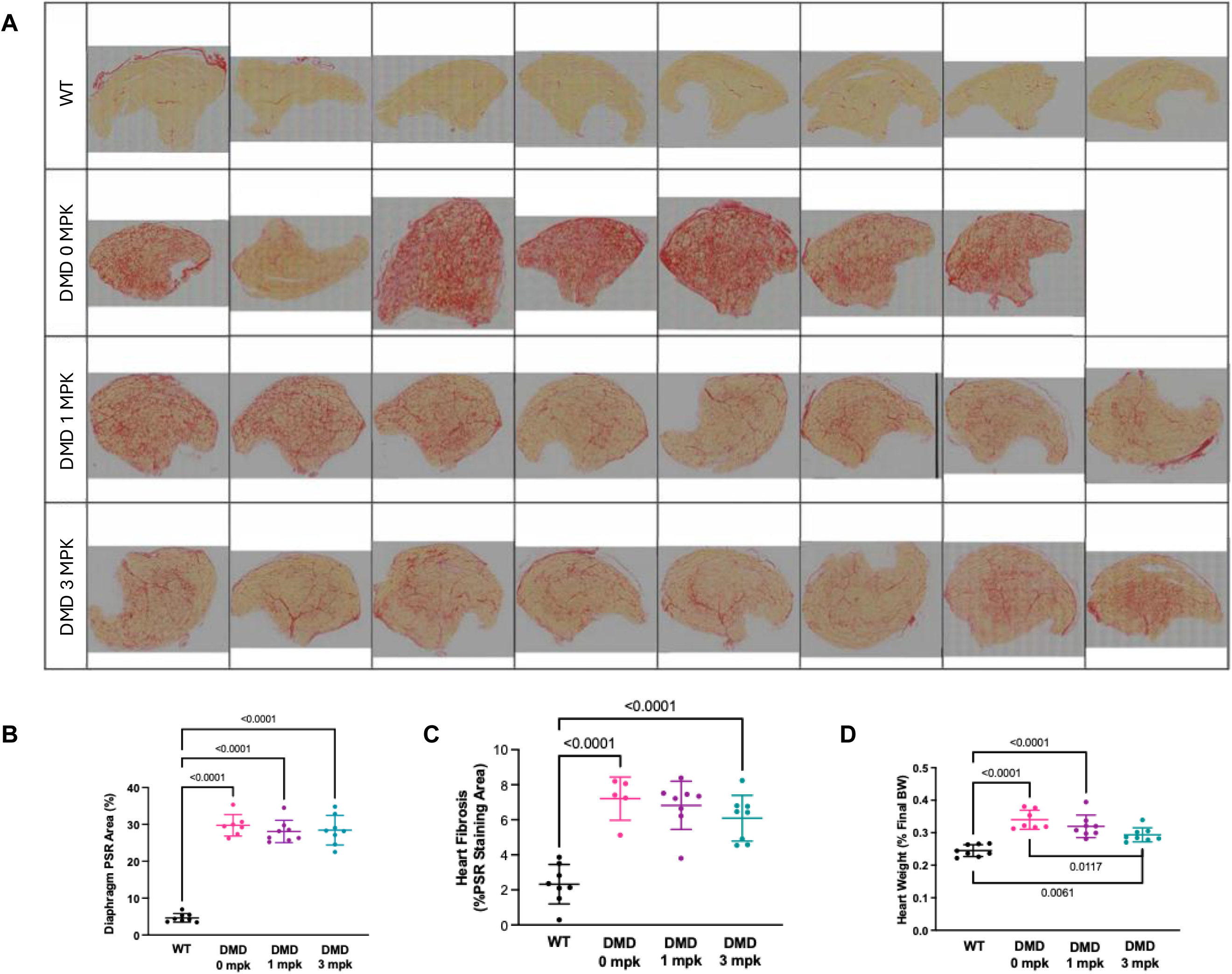
Fibrosis across the TA muscle belly, diaphragm, and heart, and cardiac remodeling in IROD-treated DMD rats. (A) Representative PSR-stained longitudinal sections spanning the entire TA muscle belly from all groups, illustrating consistency of fibrosis reduction across the full muscle extent. (B) PSR-positive area in diaphragm muscle across groups. All DMD groups exhibited significantly greater diaphragm fibrosis than WT controls, with no significant differences between treatment doses within the 8-week window. (C) PSR-positive area in cardiac muscle across groups. All DMD groups showed significantly higher cardiac fibrosis than WT controls; no significant reduction was observed with IROD treatment. (D) Relative cardiomegaly (heart weight normalized to final body weight). Treatment with 3 mg/kg IROD significantly reduced relative cardiomegaly compared with vehicle-treated DMD controls. For all dot plots, data are mean ± SEM; n = 5–8 per group. All statistical comparisons by two-way ANOVA; *P < 0.05, **P < 0.01, ***P < 0.001.

**Supplementary Fig. 4.**
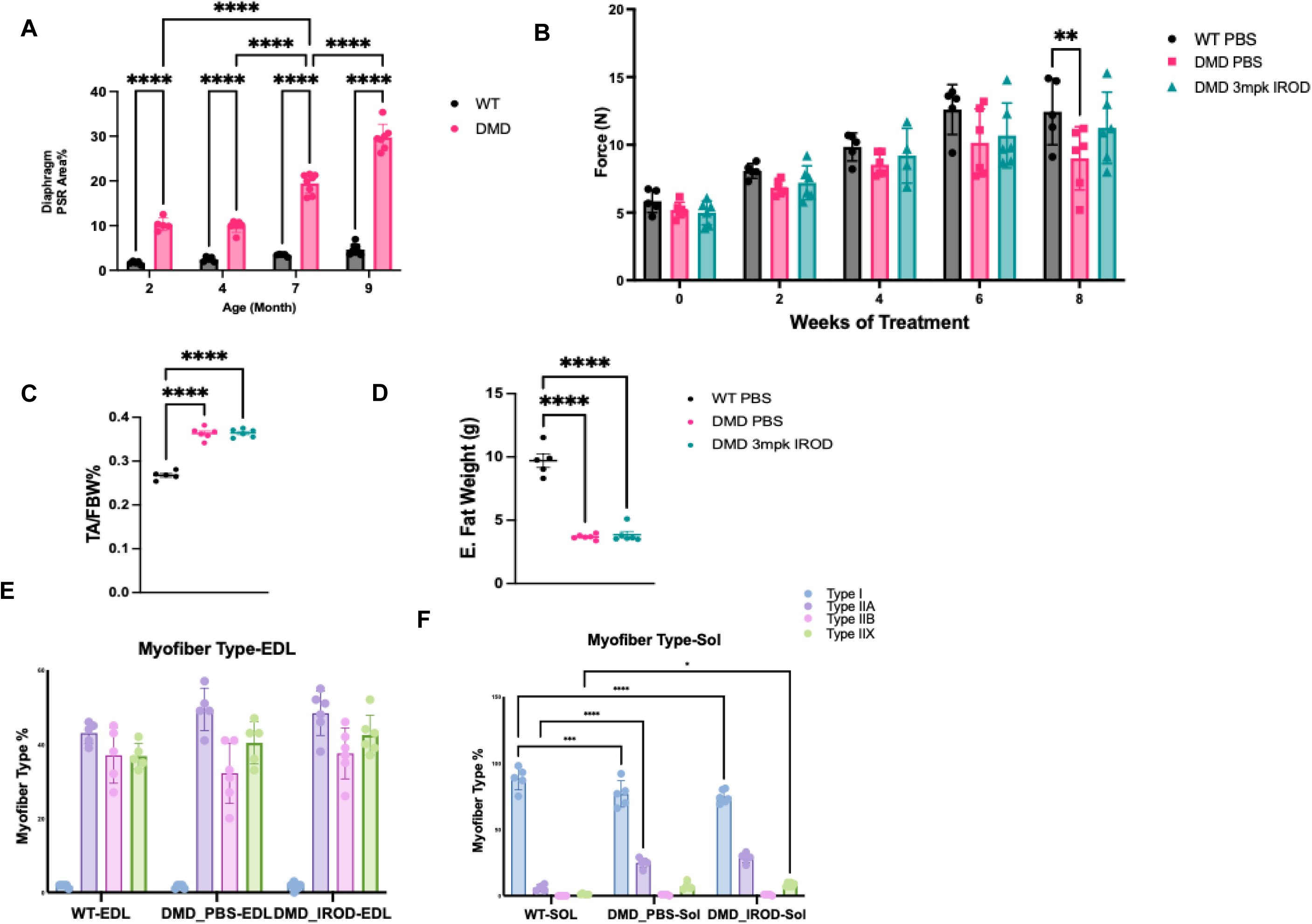
Natural history of diaphragm fibrosis, grip strength trajectory, body composition, and fiber type distribution in EDL and soleus of young IROD-treated DMD rats. (A) PSR-positive area in diaphragm of WT and untreated DMD rats at 2, 4, 7, and 9 months, establishing the natural history of diaphragm fibrosis in the DMD model. Data are mean ± SEM; n = 5–8 per group. (B) Grip strength assessed every other week over the 8-week treatment period. No significant effect of IROD on grip strength was observed. Grip strength deficits relative to WT were not detectable until 4 months of age. Data are mean ± SEM; n = 5–6 per group. (C) TA muscle weight normalized to final body weight. DMD rats exhibited a significantly higher TA/body weight ratio than WT controls, with no significant effect of IROD treatment. (D) Epididymal fat weight. DMD rats showed significantly lower epididymal fat weight than WT controls; no significant effect of IROD treatment was observed. (E) Myofiber type proportions in the EDL. No significant fiber-type changes were observed with IROD treatment. (F) Myofiber type proportions in the soleus. No significant fiber-type changes were observed with IROD treatment. For panels C–F, data are mean ± SEM; n = 5–6 per group. All statistical comparisons by two-way ANOVA; *P < 0.05, **P < 0.01, ***P < 0.001.

**Supplementary Fig. 5.**
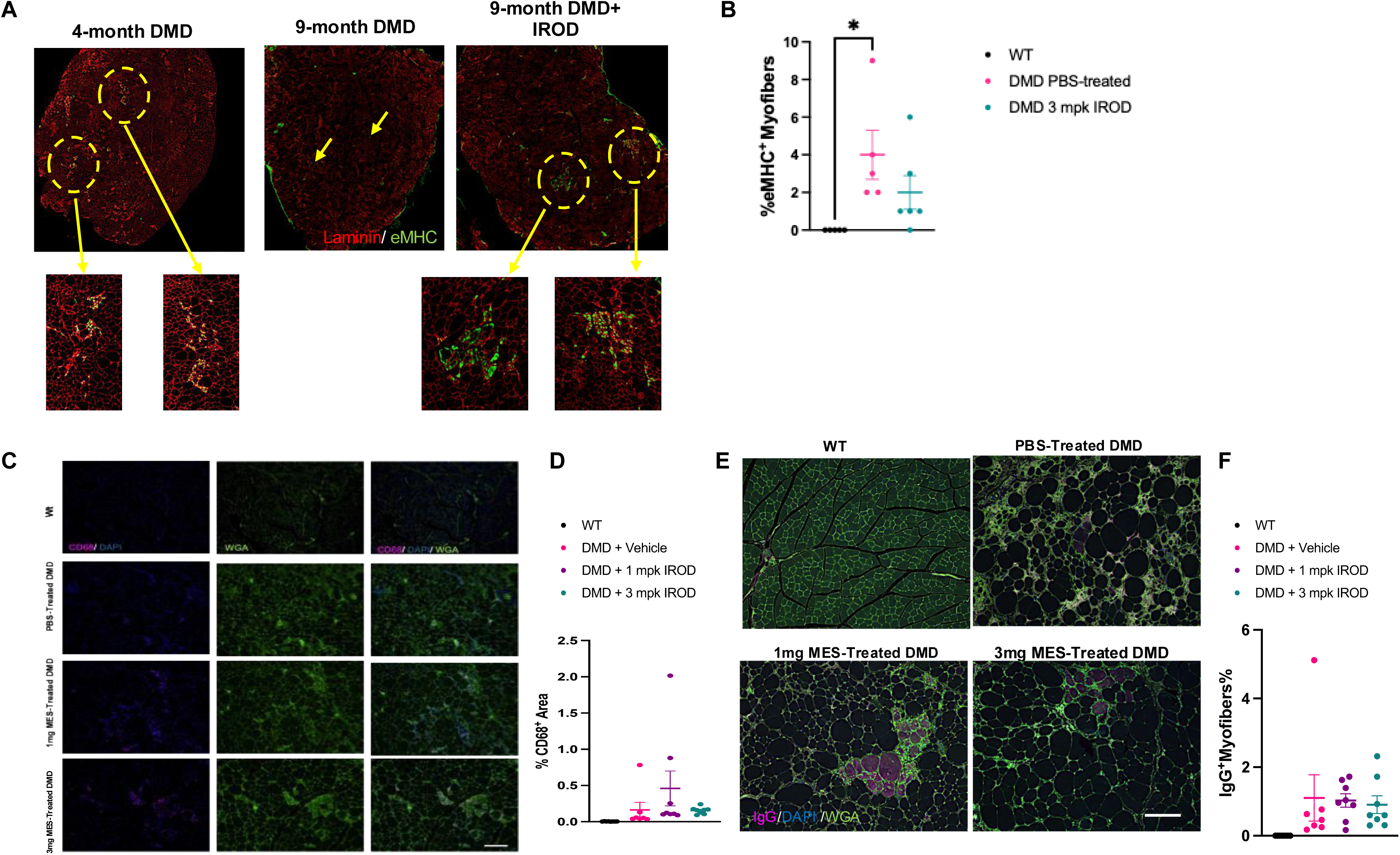
Stage-dependent eMHC+ regenerative activity and assessment of inflammation and sarcolemmal integrity in IROD-treated DMD rats. (A) Representative eMHC immunostaining in TA cross-sections from 4-month-old untreated DMD, 9-month-old untreated DMD, and 9-month-old 3 mg/kg IROD-treated DMD rats. Insets highlight clustered eMHC^+^ patterns present in 4-month-old and IROD-treated 9-month-old animals, largely absent in untreated 9-month-old DMD muscle. (B) Percentage of eMHC-positive myofibers in TA of 4-month-old DMD rats treated from 2 months with vehicle or 3 mg/kg IROD. Both groups exhibited elevated eMHC^+^ percentages with no significant difference between groups, consistent with a ceiling effect at this stage. Data are mean ± SEM; n = 4–5 per group. (C) Representative CD68 immunostaining in TA cross-sections across groups at 9 months. Scale bar = 50 μm. (D) CD68-positive area as a percentage of total TA cross-sectional area. No significant differences were observed between IROD-treated and vehicle-treated DMD animals. Data are mean ± SEM; n = 7–8 per group. (E) Representative cytoplasmic IgG immunostaining identifying myofibers undergoing active sarcolemmal disruption. No IgG-positive fibers were detected in WT rats. Scale bar = 50 μm. (F) Percentage of IgG-positive myofibers. Approximately 1% of fibers were IgG-positive across all DMD groups; IROD treatment did not significantly alter this proportion. Data are mean ± SEM; n = 7– 8 per group. All statistical comparisons by two-way ANOVA; *P < 0.05, **P < 0.01, ***P < 0.001.

## Materials and Methods

### Animals and study design

DMD rats, generated by CRISPR/Cas9-mediated frameshift mutation of the *Dmd* gene on a Wistar-Imamichi background, were used throughout this study (16). Animals were housed under standard conditions (12-hour light/dark cycle, 22 ± 2°C, 50 ± 10% relative humidity) with ad libitum access to standard rodent chow and water. All procedures were conducted at the Modified Barrier Facility (MBF) at the University of British Columbia (UBC) in accordance with the guidelines of the Canadian Council on Animal Care (CCAC) and under protocols approved by the UBC Animal Care Committee (ACC).

For the late-intervention cohort, 31 male DMD rats (approximately 7 months of age; mean body weight 460 ± 12 g) were allocated to three treatment groups (n = 7–8 per group) receiving weekly subcutaneous injections of vehicle (0 mg/kg), 1 mg/kg, or 3 mg/kg IROD for 8 weeks. Eight age-matched male WT rats served as untreated controls. For the early-intervention cohort, 17 male rats at 8 weeks of age were used: 5 WT controls and 12 DMD rats allocated to two groups (n = 6 per group) receiving weekly subcutaneous injections of vehicle or 3 mg/kg IROD for 8 weeks. For biodistribution studies, separate cohorts of 3 WT and 3 DMD male rats at 7 months of age were used for tritium-labeled IROD biodistribution and ⁹⁹ᵐTc-MDP SPECT/CT imaging, respectively. For natural history characterization of diaphragm fibrosis, 44 male rats at 2, 4, 7, and 9 months of age were used, comprising both WT and untreated DMD animals at each time point.

### Drug preparation

Irodanoprost (IROD; MES1022) and its active moiety MES1002 were provided by Dr. Robert Young (Simon Fraser University, Burnaby, BC, Canada). IROD is a mineral-targeted prodrug comprising a phosphonate-based bone-binding moiety, a hydrolyzable ester linker, and the EP4-selective prostaglandin E2 analog MES1002. The active moiety is released upon esterase-mediated hydrolysis of the linker in vivo. Both compounds were stored at −20°C until use. For in vivo administration, IROD was dissolved in phosphate-buffered saline (PBS) and administered by weekly subcutaneous injection at doses of 1 or 3 mg/kg; vehicle-treated animals received equivalent volumes of PBS. For in vitro experiments, MES1002 was dissolved in dimethyl sulfoxide (DMSO) and diluted to working concentrations in cell culture medium immediately prior to use.

### Biodistribution studies

For tritium-labeled IROD biodistribution, WT and DMD rats were transferred to the Simon Fraser University (SFU) animal facility and acclimatized for 5–6 days prior to dosing. The dosing solution was prepared to deliver 1 mg/kg with a radioactivity of 4 μCi per rat in a fixed volume of 1 mL, prepared fresh on the day of administration and injected within 1 hour by subcutaneous injection between the shoulder blades. Animals were sacrificed 24 hours post-dose and tissues were collected. Tissue samples were analyzed using an R.J. Harvey OX-300 Biological Oxidizer; samples were combusted at 800°C in a pure oxygen environment, converting organic components to tritiated water and CO₂, which were collected in scintillants and counted. Results are expressed as percentage of injected dose per gram of tissue (%ID/g).

For SPECT/CT imaging, WT and DMD rats (mean body weights 583 g and 464 g, respectively) received a single intravenous injection of 500 μL of ⁹⁹ᵐTc-MDP (injected activity approximately 90 MBq for WT and 100 MBq for DMD). SPECT imaging was performed at 5 hours post-injection using a VECTor/CT multimodal preclinical scanner (MILabs, The Netherlands) with a UHRRM rat pinhole collimator; a static acquisition of a single 30-minute frame was obtained. Animals were maintained under light isoflurane anesthesia on a heated bed throughout imaging. The ⁹⁹ᵐTc photopeak was centered at 140 keV with a 25% energy window. SPECT images were reconstructed using a pixel-ordered subset expectation maximization (POSEM) algorithm (16 subsets, 16 iterations; isotropic voxel size 0.4 mm), decay-corrected, and attenuation-corrected following CT registration. Reconstructed volumes were post-filtered with a 3D Gaussian filter for visualization. CT scans were acquired at 65 kV and 615 μA with two frames of 180 projections over 360° and reconstructed using SkyScan NRecon software (voxel size 0.169 mm³). Following imaging, rats were euthanized under deep isoflurane anesthesia, blood was collected by cardiac puncture, and organs and hindlimb muscles were dissected, weighed, and measured for radioactivity using a calibrated HIDEX gamma counter (Turku, Finland). All activities were decay-corrected to the time of injection and expressed as %ID/g.

### Behavioral and functional assessments

Animals were monitored daily throughout the study period using standard UBC clinical assessment forms. Behavioral activity was scored on a three-point scale from Grade 0 (normal activity) to Grade 2 (isolated, minimally active, and unresponsive to disturbance). Porphyrin staining — a porphyrin-pigmented ocular secretion that serves as an indicator of chronic stress or illness — was graded from 0 (no staining) to Grade 2 (extensive staining on the face, paws, and back that has not been groomed away).

Ex vivo contractile properties of the left extensor digitorum longus (EDL) muscle were assessed using the Aurora Scientific 1300A system (Aurora Scientific, Canada). Immediately following dissection, the EDL was immersed in oxygenated Ringer’s lactate solution (95% O₂/5% CO₂, pH 7.4, 30°C). Muscles were equilibrated at a resting tension of 4 mN for 10 minutes. Supramaximal stimulation conditions were established using 0.5 ms pulses to determine optimal muscle length (L₀), followed by maximum tetanic stimulation at 150 Hz to measure peak tetanic force. Active stiffness was measured by imposing a 10% length change during full tetanic contraction. Muscle cross-sectional area (CSA) was estimated as: CSA (mm²) = weight (mg) / [density (mg/mm³) × length (mm)], using a mammalian skeletal muscle density of 1.06 mg/mm³. Specific force and normalized stiffness were calculated by dividing absolute values by CSA.

Neuromuscular function was assessed by four-limb grip strength testing using a Bioseb grip strength meter (Bioseb, France). Animals were habituated to the procedure over five training sessions prior to 2 months of age. Beginning at the experimental start point, grip strength was assessed every two weeks. At each session, animals grasped a wire grid with all four limbs and were gently pulled horizontally by the tail until release; the highest of three consecutive measurements was recorded. Results are expressed in Newtons (N).

### Histology and immunostaining

Skeletal muscle tissues (tibialis anterior, EDL, soleus, and diaphragm) and cardiac muscle were harvested at study end and processed by two methods: embedding in optimal cutting temperature (OCT) compound for cryosectioning at 10 μm, or fixation in 10% neutral buffered formalin and paraffin embedding (FFPE) for sectioning at 5 μm. Sections were imaged using a Nikon fluorescence microscope.

FFPE cross-sections were stained with hematoxylin and eosin (H&E) for assessment of general muscle morphology, myofiber size distribution, and centronucleation. Minimum Feret diameter was quantified from H&E-stained sections; the coefficient of variation (CV) of Feret diameter was calculated as a measure of fiber size heterogeneity. Picro-Sirius Red (PSR) staining was performed on FFPE cross-sections and longitudinal sections to quantify interstitial fibrosis, expressed as PSR-positive area as a percentage of total muscle cross-sectional area. Both cross-sections and longitudinal sections were used to confirm consistency of fibrosis measurements across the muscle belly.

For immunofluorescence, FFPE sections underwent deparaffinization and antigen retrieval prior to antibody incubation. OCT-embedded cryosections were used for fiber-type immunostaining. Intramuscular fat accumulation was quantified using anti-Perilipin antibody (Sigma, Cat# P1873; 1:100) with goat anti-rabbit IgG Alexa Fluor 647 secondary antibody (Invitrogen, Cat# A21245; 1:500); Perilipin-positive area was expressed as a percentage of total muscle cross-sectional area. Sarcolemmal integrity was assessed by cytoplasmic IgG staining using Wheat Germ Agglutinin (Thermo Fisher Scientific, Cat# W11261; 1:500) to label fiber boundaries and goat anti-rat IgG Alexa Fluor 647 (Invitrogen, Cat# A21247; 1:500) to detect endogenous IgG infiltration in membrane-disrupted fibers. Macrophage infiltration was assessed by co-staining with anti-CD68 antibody (Abcam, Cat# ab31630; 1:200) and Wheat Germ Agglutinin (1:500), with goat anti-mouse IgG1 Alexa Fluor 647 secondary antibody (Invitrogen, Cat# A21240; 1:500); CD68-positive area was expressed as a percentage of total muscle cross-sectional area.

Myofiber type composition was assessed by multiplex immunofluorescence using the following primary antibodies: BAD5 (anti-MHC Type I; AbLab; 1:200), SC-71 (anti-MHC Type IIa; AbLab, Cat# AB0001806; 1:40), BFF3 (anti-MHC Type IIb; DSHB, 291 μg/mL; 1:10), BF-F35 (anti-MHC Type IIx; DSHB, 221 μg/mL; 1:40), F1.652 (anti-embryonic MHC; DSHB, 13 μg/mL; 1:40), and anti-Laminin (Abcam, Cat# ab11575; 1:200) to delineate fiber boundaries. Fiber type proportions were expressed as a percentage of total myofibers per section. eMHC-positive fibers were additionally assessed for clustered distribution patterns as a morphological indicator of synchronized regeneration.

Histological quantification was performed using FiberSight, an ImageJ/Fiji plugin for skeletal muscle image analysis developed at the Rossi Laboratory, University of British Columbia. FiberSight distributes fine-tuned Cellpose segmentation models optimized for skeletal muscle morphometry in both physiological and pathological tissue states. Standard generalist segmentation models perform poorly in dystrophic muscle, where fibro-fatty infiltration and non-uniform staining impair accurate fiber border delineation. To address this, baseline Cellpose models (Cyto2 and Cyto3) were fine-tuned on manually annotated image patches of mouse and rat skeletal muscle across three staining protocols — WGA, H&E, and PSR — curated by expert annotators. In tissue exhibiting fibro-fatty infiltration, fine-tuning improved average segmentation precision (IoU = 0.5) by 0.34 for PSR, 0.30 for H&E, and 0.15 for WGA compared with the generalist baseline models. FiberSight is freely available at https://github.com/ian-coccimiglio/FiberSight (*49*).

### Fibro-adipogenic progenitor isolation and differentiation assay

Fibro-adipogenic progenitors (FAPs) were isolated from tibialis anterior muscles of 7-month-old DMD rats by fluorescence-activated cell sorting (FACS). Muscles were minced and enzymatically digested, and mononuclear cells were isolated by density gradient centrifugation. FAPs were plated in growth medium and allowed to adhere overnight before treatment. To induce fibrogenic differentiation, cells were treated with TGF-β (5 ng/mL) for 72 hours in the presence or absence of MES1002 at 125, 250, or 500 μM. Fibrogenic differentiation was assessed by immunofluorescence staining for α-smooth muscle actin (α-SMA) and collagen I. α-SMA-positive area was quantified as a percentage of total cell area per field.

### RNA sequencing and transcriptomic analysis

Total RNA was extracted from tibialis anterior muscles harvested one week after the final dose. Initial alignment was performed using Illumina’s DRAGEN pipeline by the UBC Biomedical Research Centre Sequencing Core, returning Salmon-aligned gene transcripts. Transcript-level estimates were converted to gene-level counts using tximport (v1.32.0) with countsFromAbundance set to ‘LengthScaledTPM’, using *Rattus norvegicus* genome annotations (Rnor_6.0.97.gtf) (*50*). Genes with fewer than 5 counts in at least 5 of 15 samples were excluded prior to analysis. Differential expression analysis was performed in R (v4.4.0) using DESeq2 (v1.44.0) with Wald’s test(*51, 52*), comparing vehicle-treated DMD versus WT controls and IROD-treated versus vehicle-treated DMD rats. Genes with an absolute log₂ fold change > 1 and adjusted P-value < 0.05 (Benjamini–Hochberg correction) were considered differentially expressed. Principal component analysis was performed on variance-stabilizing transformed (VST) counts. Gene Ontology biological process enrichment analysis was performed using clusterProfiler (v4.12.0) with the enrichGO function(*53*), and redundant GO terms were removed using the simplify function from the DOSE package (v3.30.0)(*54*). Fold enrichment was calculated as the ratio of input genes to background genes within each GO term. Heatmap visualizations were generated using row-normalized Z-scores of the top 20 differentially expressed genes per contrast, sorted by adjusted P-value.

### Statistical analysis

Data are presented as mean ± standard deviation (SD) unless otherwise stated in figure legends. Biodistribution data are presented as mean ± standard error of the mean (SEM). Statistical analyses were performed using GraphPad Prism (Version 10; GraphPad Software, San Diego, CA, USA). A significance threshold of P < 0.05 was applied throughout. Body weight trajectory data were analyzed by two-way ANOVA with Dunnett’s post-hoc test for multiple comparisons against the vehicle-treated DMD group. All other group comparisons were performed by two-way ANOVA with Tukey’s post-hoc test. For RNA-seq data, differential expression analysis and statistical testing were performed as described above using DESeq2 with Benjamini–Hochberg correction for multiple comparisons.

